# Unifying constraints linking protein folding and native dynamics decoded from AlphaFold

**DOI:** 10.64898/2026.01.08.698426

**Authors:** Zecheng Zhang, Weitong Ren, Liangxu Xie, Yuxiang Zheng, Xingyue Guan, Jun Wang, Wenfei Li, Qian-Yuan Tang

## Abstract

The interplay between protein folding and native dynamics remains a central question in biophysics. Analyzing an extensive set of AlphaFold-predicted structures, we uncover a robust relationship between folding topology (contact order) and native dynamics (fluctuation entropy), showing that long-range contacts that slow folding also restrict conformational flexibility across protein sizes and taxonomic groups. Scaling analysis reveals that this relationship, together with its chain-length dependence, is consistent with power-law–like trends, reflecting common organizing constraints of protein architecture. Across species, increasing organismal complexity is associated with proteome-wide shifts toward lower contact order and higher fluctuation entropy. Together, evidence from folding, stability, and functional dynamics converges on unifying constraints, revealing an intrinsic physical organizing principle captured by AI models.

---

The interplay between protein folding and function presents a central biophysical challenge in deciphering protein evolution. Folding governs the acquisition of a polypeptide’s functional three-dimensional structure (microseconds to seconds) [1–3], while native dynamics—the ensemble of post-folding conformational fluctuations (nanoseconds to microseconds) [4, 5]—enable catalysis, binding, and allosteric regulation. These processes are interconnected through the protein’s energy landscape: the same interactions that guide folding define the topology and barriers constraining functional motions. Quantifying this relationship between structural organization and dynamic behavior is essential for decoding the physical principles driving protein evolution.

Recent advances in artificial intelligence (AI) have revolutionized structural biology, providing new tools to address these fundamental questions. Tools such as AlphaFold 2 (AF2) [6], RosettaFold [7], and ESMFold [8] now produce near-atomic resolution predictions, while the AlphaFold Database (AFDB) contains hundreds of millions of structures across diverse organisms [9, 10]. This breakthrough enables systematic exploration of the connection between protein folding and function across evolutionary lineages, overcoming the severe sampling limitations of experimentally determined structures. Yet, despite this progress, we lack general physical laws describing how structural properties and dynamical flexibility scale with protein size and organismal complexity. In this Letter, we perform a large-scale analysis of AIpredicted protein structures and identify unifying constraints linking folding topology to native-state dynamics. Analyzing hundreds of thousands of AF2-predicted structures from 45 diverse organisms, we identify a robust negative correlation between contact order (a measure of folding complexity) and fluctuation entropy (quantifying native-state flexibility). Scaling analysis shows that this inverse relationship and its chain-length dependence are consistent with simple power-law–like trends, reflecting underlying structural constraints. Comparing proteins of similar chain lengths across organisms, we find that those from more complex species fold with fewer longrange contacts and exhibit greater native-state fluctuations, revealing a structural signature of proteome organization shaped by functional and evolutionary demands. Independent evidence from sequence entropy inferred from protein language models [8] and melting-temperature data [11] supports this picture, indicating that protein evolvability and stability are constrained by topology-dependent flexibility. Taken together, these findings point to physical connections among folding topology, dynamic flexibility, and evolutionary diversity. These connections reflect intrinsic properties of natural proteins, are captured by AI-based structure predictions, and made explicit through physical analysis.

## Basic Setups

We introduce two key metrics used in this study. The first is *contact order* (CO), which quantifies the non-locality of residue contacts in a protein’s native structure [12–14]. CO is defined as:

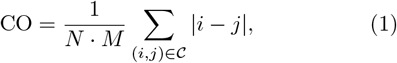

where N is the chain length, 𝒞 is the set of residue pairs with C_*α*_ distance r_*ij*_ < r_CO_ = 8 A and sequence separation |I − j| ≥ 4 (i.e., long-range contacts; see Fig. 1A), and M = |𝒞| is their total number. CO represents the average sequence distance of long-range contacts, normalized by N. Theoretical, computational, and experimental studies showed that proteins with lower CO—entailing lower entropic cost and shorter search times in conformational space—tend to fold faster [15–19].

**FIG. 1.**
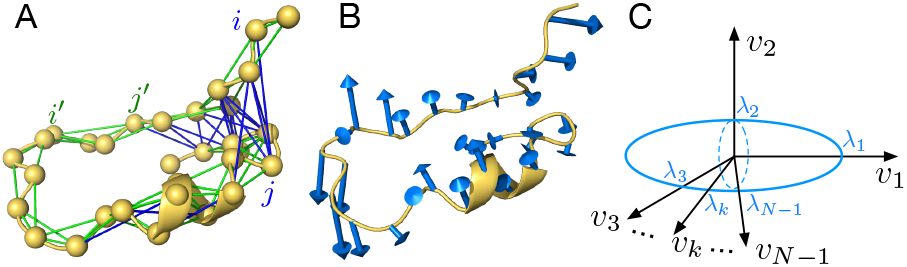
Schematic illustration of contact order and fluctuation entropy. (A) Elastic network model (ENM) of an example protein (UniProt ID: A0A0R0FF49). Nodes represent C_*α*_ atoms of residues; edges represent harmonic interactions between residues. Long-range contacts (|*i− j*| *≥* 4, blue) and short-range interactions (|*i−j*| *<* 4, green) define folding topology. (B) The first normal mode predicted by ENM, illustrating collective motions of residues around the native conformation. (C) Fluctuation entropy *S* quantifies the logarithm of accessible conformational space volume around the native structure, depicted as a high-dimensional ellipsoid with principal axes corresponding to normal mode amplitudes.

The second metric, the *fluctuation entropy* (S), serves as a proxy for native-state structural dynamics by quantifying the effective volume of conformational space accessible around the native structure. [20–22]. In the Elastic Network Model (ENM), the Hessian matrix eigenmodes describe collective motions of the structure [5, 23, 24]. Each mode contributes a variance λ_*k*_ to the covariance matrix C (with entries 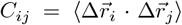), such that S = log det C = Σ _*k*_ log λ_*k*_ aggregates the contributions from all modes (Fig. 1B). Higher S values indicate greater collective fluctuations, whereas lower values reflect restricted conformational motions. To complement this linearized description, we introduce the *sequence entropy* H as an evolutionary measure of tolerated structural variation. Highly variable sites tend to permit local flexibility, whereas conserved sites reflect mechanical constraints [22, 25]. Because mutational tolerance integrates nonlinear physical effects, H serves as a data-driven proxy for structural adaptability, capturing features beyond the linear ENM and independent of AF2 modeling (see Appendix A).

This analysis is based on AF2-predicted structures for 45 organisms from AFDB (SI, Table S1), retaining reliable models with mean pLDDT > 70 [26]; results are robust across thresholds. While experimentally determined structures remain invaluable, they may be biased toward proteins that crystallize or express readily [27], whereas AI-predicted models enable proteome-wide coverage and a broader view of structure–function relationships. AF2 predictions generally achieve near–atomic resolution [28, 29] and reliably reproduce residue–residue contacts, providing sufficient accuracy for CO and S calculations. Although AF2 does not explicitly represent full conformational ensembles, its predictions nevertheless encode key signatures of the energy landscape and mutational effects [30–32]. Additional validation analyses are provided in Appendix B.

### Statistical and Theoretical Analysis

To uncover the link between protein folding and native dynamics, we analyzed proteins with similar chain lengths (225 ≤ N < 275) from 16 model organisms on the CO–S plane (Fig. 2A). Averaging S within CO bins reveals a clear negative correlation. Given that lower CO predicts faster folding [12–14, 18], this trend implies that rapidly folding proteins tend to exhibit larger native-state fluctuations and softer collective modes. Analysis across diverse organisms, including archaea (*M. jannaschii*), bacteria (*E. coli*), unicellular eukaryotes (*S. cerevisiae*), multicellular invertebrates (*C. elegans*), and humans (*H. sapiens*), demonstrates the broad consistency of the CO–S relationship, indicating common physical constraints rather than organism-specific evolutionary adaptations. Notably, this relationship is also encoded at the sequence level. For the same protein set used in panel A, grouping by sequence entropy H reveals that CO decreases and S increases with H (Fig. 2B,C), with fulldata and median fits nearly overlapping, confirming the robustness of the trend. Because this analysis uses only sequence-derived H, independent of ENM-based S yet consistent with the AF2-derived trend, the CO–S connection holds beyond the linear regime. Together, these results link folding topology, native dynamics, and evolutionary variability.

**FIG. 2.**
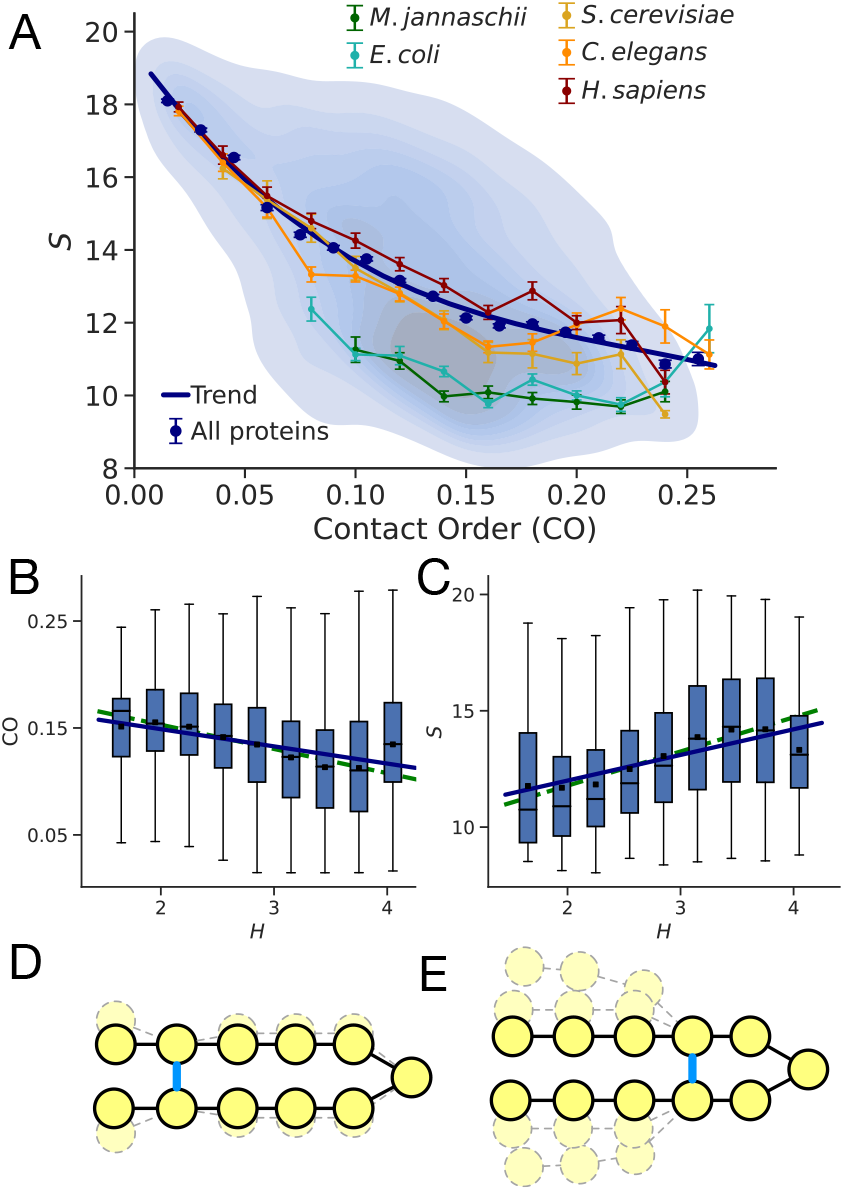
Contact order (CO) negatively correlates with fluctuation entropy *S*. (A) Heat map of *S* versus CO for proteins of similar chain length (225*≤ N <* 275) across 16 model organisms, with the averaged trend and selected organism curves highlighted. (B–C) Box plots of CO and *S* grouped by sequence entropy *H* (outliers omitted). For both quantities, binned means (dots with solid-line fits) and fits to the full data (dashed lines) closely coincide, indicating consistent negative CO–*H* and positive *S*–*H* correlations. (D–E) Schematic illustrating the effect of residue sequence distance on *S*: (D) a non-local contact significantly lowers *S*, whereas (E) a local contact preserves higher *S*.

The conceptual model in Figs. 2D,E explains the mechanism: with uniform edge weights, each added contact contributes similarly to energetic stabilization, but its entropic effect depends on sequence separation. Longrange contacts impose stronger geometric constraints and reduce fluctuations (panel D), whereas local contacts preserve greater conformational freedom (panel E). This is consistent with folding studies showing contact-induced entropy loss during structure formation [33–36], with evidence that contact patterns regulate native-state fluctuations [37, 38], and with recent AI-based studies indicating a local-first, global-later folding hierarchy [39]. Here, we derive the CO–S relationship directly from the global organization of the residue contact network. From a graph-theoretic perspective, higher S corresponds to a reduced number of spanning trees in the contact graph (SI, Sec. V). Thus, reducing long-range contacts while reinforcing local ones lowers the number of spanning trees, stabilizing hydrophobic cores while maintaining flexibility, consistent with our earlier finding that protein structures balance robustness and plasticity [21]. Additional analyses (SI, Sec. IV C) show that long-range contacts provide the dominant topological constraints of the fold, and that their disruption introduces anharmonic effects and triggers larger-scale motions.

Remarkably, different functional classes exhibit systematic shifts along the CO–S plane: regulatory proteins favor lower CO and higher S, whereas metabolic enzymes show the reverse pattern (SI, Sec. IV D, Fig. S10), reflecting function-dependent tuning of topology and flexibility. The trend further extends to other structural descriptors (Appendix C), indicating a broader topology–dynamics relationship beyond the linear ENM regime. Consistently, experimental thermal-stability data show the same directionality [11]: in human proteins, higher CO and lower S correlate with higher melting temperatures T_*m*_ (Appendix D), validating the physical relevance of CO and S in shaping protein stability.

### Scaling Analysis

Having established the CO–S correlation at fixed chain length N, we next examine how protein size modulates this relationship. We therefore perform a scaling analysis to quantify how slow-mode contributions to S depend jointly on CO and N. This approach is motivated by prior work showing that native proteins exhibit dynamical signatures of criticality [40– 47]. Focusing on the dominant collective component, we analyze the first normal mode, whose entropic contribution is log λ_1_. Grouping proteins by chain length N, we observe a consistent negative correlation between CO and λ_1_ (Fig. 3A), consistent with a phenomenological powerlaw–like scaling, λ_1_ ~CO^*−α*^ with *α* = 1.027, given the limited range of CO. We next examine how λ_1_ scales with N. As shown in Fig. 3B, λ_1_ ~*N*^*β*^ with β = 1.540. To isolate the effect of N, we fix CO and find that λ_1_ still increases with N within fixed-CO groups (Fig. 3C), following λ_1_ ~N^*γ*^, with γ = 0.791. By combining the scaling relations, we derive the composite scaling λ_1_ ~CO^*−ζ*^, with ζ = *αβ*/(*β* −γ) ≈ 2.11. This prediction closely matches the direct fit to data (ζ = 2.293; Fig. 3D).

**FIG. 3.**
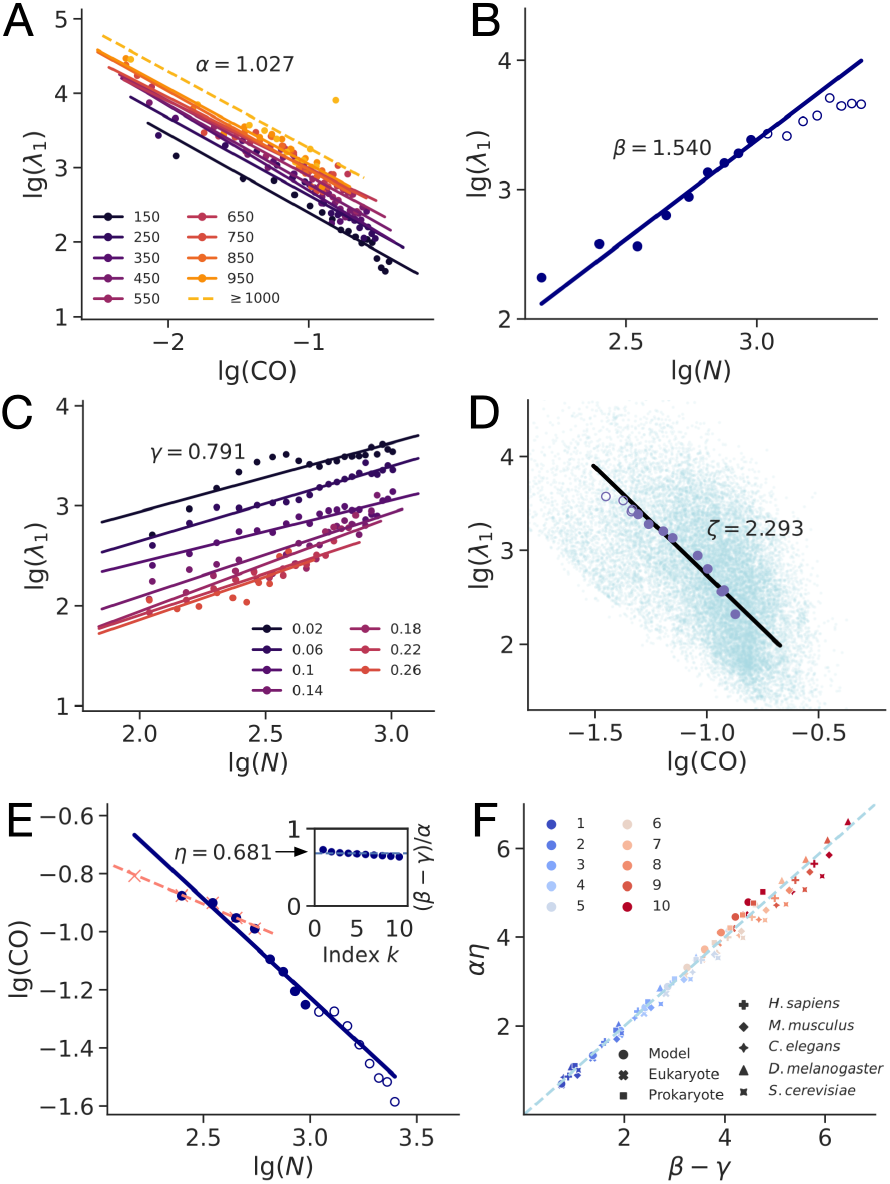
Scaling Analysis. (A) A power-law–like dependence *λ*_1_ *~*CO^*−α*^ with *α* = 1.027 is observed across different chain length groups. The group with *N* = 150 includes proteins where 100 *≤ N <* 200; similar binning applies to other ranges. (B) Fitting for proteins shows *λ*_1_*~ N*^*β*^ with *β* = 1.540. (C) Similar scaling *λ*_1_ *~ N*^*γ*^, with *γ* = 0.791, is found across different CO groups. The group with CO = 0.02 includes proteins where 0*≤* CO *<* 0.04; similar binning applies to other ranges. (D) Validation of the derived relation *λ*_1_ *~*CO^*−ζ*^. Each light blue dot represents a protein. Proteins are binned by their chain length *N*, and the average *λ*_1_ and CO values for each bin are used for the log–log fit. (E) Fitting of CO vs. *N* yields CO *~ N* ^*−η*^ with *η* = 0.681, and a separate fit for proteins with 100 *≤ N <* 600 yields *η*′ = 0.312 (pink dashed line). The inset confirms the consistency of *η* = (*β − γ*)*/α* across the first 1–10 leading modes. (F) Verification of the identity *αη* = *β − γ* across proteins from different organisms with varying numbers of included modes. All fittings are based on proteins with 100 *≤ N <* 1000, and we use proteins with *N ≥* 1000 as validation sets. Panel A uses solid and dashed lines, while panels B, D, and E use solid and empty circles to indicate fits and validations, respectively.

Our scaling analysis reveals that CO decreases with chain length as CO ~ N^*−η*^, with a fitted exponent η = 0.681 (Fig. 3E). Notably, this fit remains valid for large proteins (N ≥ 1000), supporting its applicability across the full size range. To better capture this global trend, a small-N outlier was excluded from the fit. When restricted to smaller proteins (N < 600; pink dashed line), our analysis reproduces the previously reported exponent η′≈ 1/3 [14, 48]. The steeper scaling observed over the full range suggests that longer proteins evolve to more aggressively reduce long-range contacts—likely by adopting modular or domain-like architectures that enable quasi-independent folding. Such organization minimizes topological frustration and enhances folding efficiency, particularly in large, complex proteomes.

From the scaling relations discussed above, we derive the identity *αη* = *β*− *γ*, which links the corresponding exponents (Appendix E). Control analyses, including sequence redundancy reduction, alternative binning schemes, and stratification by secondary-structure composition, show that this relation is robust, with only modest variations in exponent values and preserved internal consistency (SI, Sec. VII). The identity also holds across species and when incorporating multiple slow modes, with exponents recalculated using cumulative products 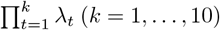 remaining consistent (Fig. 3E, inset; Fig. 3F). Taken together, these observations suggest shared constraints linking protein folding and native dynamics, reflecting an organizing pattern observed across a wide range of protein sizes and phylogenetic groups. Additional support comes from a renormalization analysis (SI, Sec. VII E), in which coarse-grained and fragmented protein structures also obey a similar scaling relation, indicating resolution invariance of the underlying physical constraint. Such scale invariance is characteristic of critical phenomena, suggesting a physical rather than purely biological origin of protein architecture and motions. This behavior is also informed by polymer physics (SI, Sec. VIII), where loop-closure statistics provide a natural baseline for the fitted scaling exponents and complement evolutionary and functional interpretations.

### Cross-organism comparisons

We next examine how protein folding and native dynamics vary across species by comparing proteins of similar chain length N and analyzing organism-level distributions of CO and S. To relate these patterns to organismal complexity, we use the total number of proteins and the cumulative chain length of the proteome as proxies. Although the definition of biological complexity remains debated [49], several studies have shown strong correlations among genome size, number of cell types, and proteome size [50–52], supporting the validity of our chosen metrics. This enables a quantitative, cross-organism comparison of protein structure and dynamics in relation to biological complexity.

As shown in Fig. 4A, proteins of similar chain length exhibit an inverse relationship between mean CO and organismal complexity, with more complex species tending toward lower CO. This trend is robust across contact definitions, chain-length windows, and filtering thresholds (Sec. VI, SI). Likewise, fluctuation entropy S correspondingly increases with organismal complexity (Fig. 4B), and both correlations persist under alternative parameters and after excluding low-confidence models (mean pLDDT < 70). Examining CO and S distributions in five representative organisms matched for chain length N (Fig. 4C–D) confirms that proteins from more complex species shift toward lower CO and higher S, consistent with increased structural modularity and flexibility [53], revealing a systematic trend in which higher organismal complexity corresponds to fewer long-range contacts and greater conformational flexibility.

**FIG. 4.**
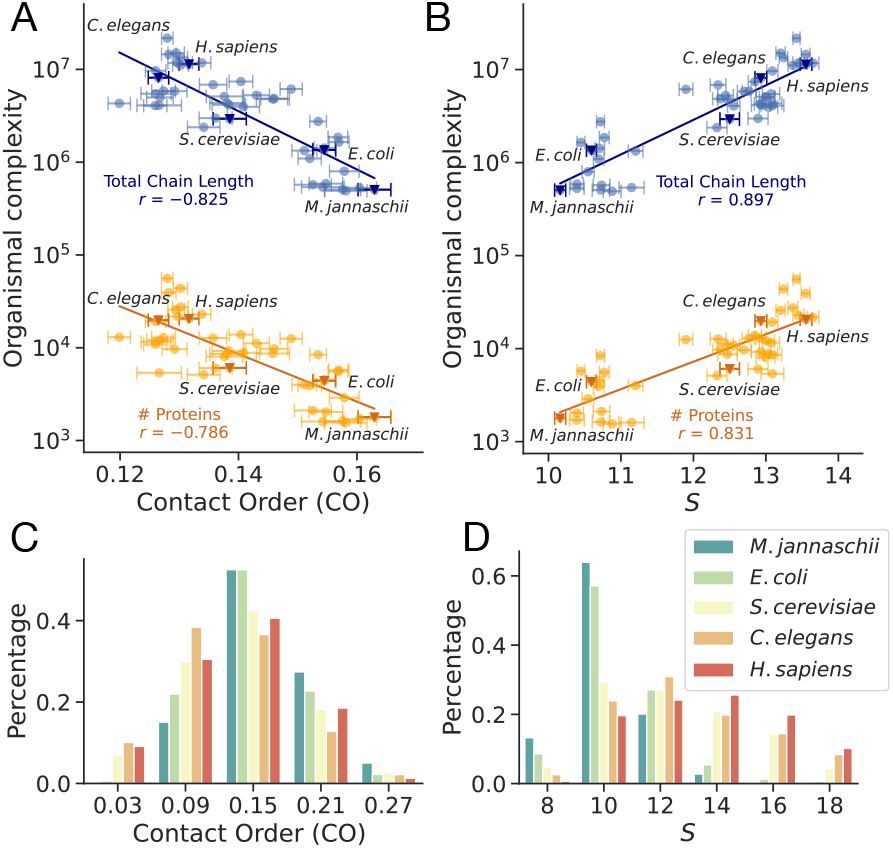
Cross-organism comparisons of CO and *S* for proteins with similar chain lengths (225 *≤ N <* 275) across 45 species. Scatter plots show correlations between organismal complexity and (A) mean CO and (B) mean *S*, with dots representing group averages and error bars indicating standard error. In panels A and B, different colors denote alternative proxies for organismal complexity. Panels C and D show the distributions of CO and *S*, respectively, for proteins from five representative model organisms.

This trend reflects an evolutionary alignment between proteome architecture and folding dynamics. As proteomes increase in complexity, with greater sequence diversity, longer proteins, and more intricate interaction networks, maintaining efficient and reliable folding becomes increasingly important in crowded cellular environments [54]. Lower CO reduces topological frustration, mitigates misfolding and aggregation risks [55], and decreases reliance on chaperone-assisted folding [56]. Such architectures also facilitate cotranslational folding [57, 58], allowing contacts to form during synthesis and supporting timely maturation. At the same time, higher S enables functional sensitivity [21, 59, 60], adaptive specialization [61, 62], and robustness to mutation [22, 63, 64]. Thus, chain length, CO, and S co-evolve to balance folding stability, adaptability, and functional precision in complex organisms.

In summary, this study uses AI-predicted protein structures to uncover a topology-controlled scaling relation between folding architecture and native-state dynamics across species and system sizes. Supported by correlations with sequence entropy, melting temperatures, and structural measures, this relationship shows that protein stability and evolvability are shaped by topological constraints on native-state flexibility, linking evolutionary variation, folding architecture, and functional dynamics within a unified physical framework. This framework enables efficient topology-driven protein design, including rapid estimation of mutational effects, screening of scaffolds for targeted dynamical properties, and assessment of structural designability [22]. It also supports functional analyses, such as identifying allosteric contacts via entropy changes and refining coarsegrained models. More broadly, this work illustrates how physical laws can be inferred from large-scale AI structural predictions without access to internal model parameters, extending AI-based analysis beyond static structure to function and evolution. Looking ahead, integrating this framework with models of sequence variation [32, 65], protein–protein interactions [66, 67], and functional constraints (e.g., allosteric transitions and ligand binding) [68–70] may enable proteome-scale predictive biology and advance rational protein design.

The authors thank Xiangze Zeng, Liang Tian, Lei-Han Tang and Dante R. Chialvo for their insightful discussions. This research was supported by Research Grants Council of Hong Kong (No. 22302723), Natural Science Foundation of China (Grant Nos. 12305052, 12574224, 12347102, and 22003020), Hong Kong Baptist University’s funding support (RC-FNRAIG/22-23/SCI/03), Basic Research Program of Jiangsu Province (BK20253050), and the grant of Wenzhou Institute, University of Chinese Academy of Sciences (WIUCASQD2023015).

## APPENDIX

### A. Entropic Measures

The fluctuation entropy S is computed using the Elastic Network Model (ENM). Each protein structure is coarse-grained by representing each residue with its C_*α*_ atom, and harmonic springs are assigned between residue pairs within an 8 °A cutoff. A uniform spring constant is used to construct the Hessian matrix, whose diagonalization yields the normal modes. The covariance matrix C is obtained as the pseudoinverse of the Hessian, where the inverse eigenvalue λ_*k*_ = 1/σ_*k*_ gives the variance of positional fluctuations for mode k, and σ_*k*_ is the corresponding Hessian eigenvalue (after removing the zero modes). Here we retain the eight lowest-frequency nonzero modes, which dominate collective, topology-controlled fluctuations and serve as a robust proxy for global conformational entropy. Our results are insensitive to the precise number of modes retained.

Sequence entropy H provides a data-driven measure of evolutionary variability and is computed using the pretrained protein language model `esm2_t30_150M_UR50D`, which infers site-specific amino acid distributions independently of any structural modeling [8] For a protein of length N, at each site i the PLM estimates p_*i*_(a context) by masking that position and computing the conditional distribution over the 20 amino acids (renormalized to unit mass). The site entropy is 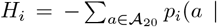 context) log_2_ p_*i*_(a | context), and the overall sequence entropy is the positional average 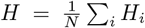. Thus, H quantifies tolerated evolutionary variability along the sequence and serves as a data-driven and nonlinear counterpart to the structural fluctuation entropy S; together, S and H reveal a unified entropic framework linking protein topology, dynamics, and evolution. We additionally confirm that the CO–H and S–H relations persist across proteins of different sizes (SI, Sec. III B).

### B. Validity of AF2-Predicted Structures for Statistical Analyses

AF2 predictions are reported to be highly precise [28, 29] and capture signatures of the underlying energy landscape, including mutational responses [30–32]. This capability arises from evolutionary grounding: multiple sequence alignments encode coevolving residue pairs that specify native contact topology rather than exact atomic detail [6, 71–73]. Complementary evidence from protein language models further shows that sequence variation patterns reflect flexibility and energetic constraints [8], reinforcing that topology and dynamics are jointly encoded at both sequence and structure levels.

AF2 models nonetheless exhibit known limitations [74]. Side-chain placement is less accurate than the backbone [75], yet it is backbone contact topology that provides the dominant constraints on folding pathways and collective motions [5, 15, 76, 77]. AF2 may underresolve proteins with multiple conformations [78], but predicted structures commonly lie along natural apo–holo transition pathways (SI, Sec. II). ENM-based fluctuation profiles remain stable across AF2 uncertainty ranges (SI, Sec. III D), supporting the use of AF2 structures for topology-governed dynamical analysis. Finally, statistical averaging across large and diverse datasets mitigates structure-specific noise, yielding robust and reproducible physical relationships. These limitations therefore do not affect the contact-topology–based quantities discussed in this paper or its main conclusions.

### C. Topology–Dynamics Relationships across Structural Metrics

Beyond the CO–S relation, similar topology–dynamics correlations emerge across other structural descriptors. One example is the modularity Q of the residue contact network. Statistical analysis shows that Q correlates negatively with CO and positively with S, indicating that fewer long-range contacts promote modular organization and enhance structural flexibility. Importantly, structural modules need not correspond to contiguous sequence segments; a single module may arise from discontinuous regions of the chain, though CO–Q correlations suggest that many real proteins fold into coherent sequence–structural sectors. Another descriptor is the fractal dimension d_*f*_ of residue packing, which correlates positively with CO and negatively with S, consistent with denser, globally connected structures constraining fluctuations. Together, these results demonstrate that the topology–dynamics relationship persists across metrics, extending beyond fluctuation entropy and the linear ENM regime (SI, Sec. VI).

### D. Thermal Stability Validation Using Meltome Atlas Data

We further validated the CO–S trend using human melting temperatures T_*m*_ from the Meltome Atlas [11]. For proteins of similar chain lengths, CO increases and S decreases with T_*m*_ (Fig. 5), with fulldata and median fits nearly overlapping, confirming the robustness of the correlation. This agreement with experimental thermal stability data supports CO and S as physically grounded descriptors of folding topology and native-state flexibility. Proteins with high CO better resist thermal unfolding, while those with greater native-state flexibility (high S) melt at lower temperatures. Thus, CO and S together provide a compact quantitative link between folding topology, dynamics, and thermodynamic stability. Further details and validations for proteins of other sizes are provided in SI Sec. VI B.

**FIG. 5.**
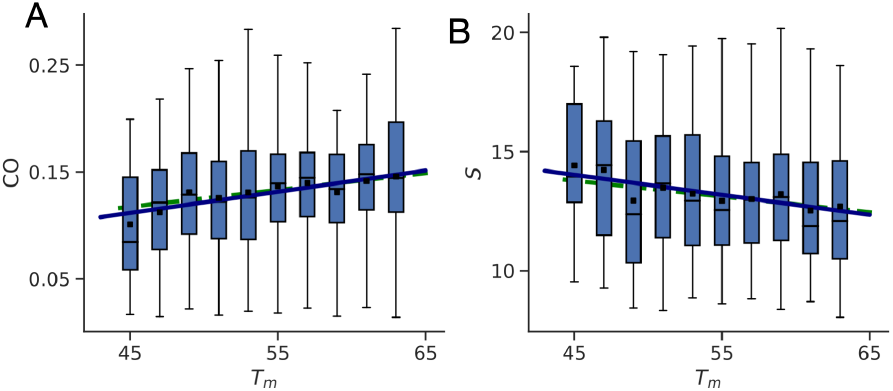
Thermal stability validation of the CO–*S* relationship. For human proteins of similar chain lengths (200 *≤ N <* 300), box plots (outliers not shown) show that (A) CO increases with melting temperature *T*_*m*_, while (B) the corresponding *S* decreases with *T*_*m*_. In both panels, fits to the full data (dashed lines) and to the binned means (dots with solid-line fits) nearly overlap, indicating consistent trends.

### E. Composite Scaling Relations

Beyond the individual scaling relations identified above, one can derive a composite relation that links them in a self-consistent manner. Here we summarize the key scaling relations introduced in the Main Text and show how they connect. Assuming a separable scaling form for the first normal-mode frequency λ_1_ as a function of chain length N and contact order (CO), we write λ_1_ ~*N*^*γ*^ CO^*−α*^, where α and γ are effective exponents obtained from fixed-N and fixed-CO subsets, respectively. From the global scaling trend we also have λ_1_ ~*N*^*β*^. Equating these two expressions gives *N*^*β*^ ~*N*^*γ*^ CO^*−α*^, so that CO^*−α*^ ~*N*^*β−γ*^. Using the empirical relation CO ~*N*^*−η*^, this becomes (*N*^*−η*^)^*−α*^ = *N*^*αη*^ ~*N*^*β−γ*^. Matching exponents yields the composite relation

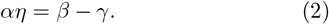

Equation 2 links the topology–size exponent η with the dynamics–size–topology exponents α, β, and γ, and implies the composite scaling λ_1_ ~CO^*−ζ*^ with ζ = αβ/(β − γ). This relation, reminiscent of scaling relations in critical phenomena, serves as an internal consistency check: independently fitted exponents are compatible with it within uncertainties, and the derived composite scaling for ζ is internally consistent across our empirical fits.

