## Supplementary Materials for "Unifying constraints linking protein folding and native dynamics decoded from AlphaFold"

### CONTENTS

I. Data and Code Availability 1
A. Dataset 1
B. Code Availability 1

II. Validity of AlphaFold-Predicted Structures for Statistical Analysis 2

III. Structure- and Sequence-based Entropy Measures 3
A. Fluctuation Entropy from Gaussian Network Model (GNM) 3
B. Sequence Entropy from Pretrained Protein Language Model 4
C. Robustness of Contact Order and Fluctuation Entropy to Parameter Variations 5
D. Consistency between Normal Mode Analysis and AlphaFold uncertainty measures 5
E. Extension to Anisotropic Network Model (ANM) 6

IV. Structural Basis of the CO–$S$ Relationship with Dynamical and Functional Implications 6
A. Representative Protein Examples Illustrating the CO–$S$ Relationship 6
B. Contact density as a source of scatter in the CO–$S$ relation 7
C. Anharmonic protein motions constrained by long-range contacts 7
D. Correlation Between the CO–$S$ Relationship and Biological Function 8

V. A Graph-Theoretical Perspective on the CO–$S$ Relationship 9
A. Intuitive Picture 9
B. Formal Proof 9
C. Extension to Weighted Graph Laplacians 11
D. Spectral–topological Interpretation of the Critical Behaviors 11

VI. Topology–Dynamics Relationships Beyond the ENM Approximation 12
A. Structural Topology Measures: Modularity and Fractal Dimension 12
B. Consistency with Experimental Melting Temperatures 13

VII. The Robustness of Scaling Analysis and Renormalization 13
A. Robustness to sequence redundancy 13
B. Data binning and fitting robustness 14
C. Secondary-structure composition 14
D. Robustness of Scaling Across Organisms and Mode Numbers 15
E. Renormalization Analysis 16

VIII. Polymer-Physics Origin of Scaling Exponents 17

IX. Robustness of Cross-Organism Analysis 18

References 19

*

### I. DATA AND CODE AVAILABILITY

#### A. Dataset

In this study, we utilized predicted structures from the AlphaFold Protein Structure Database (AFDB, version 4) for our statistical analyses. We selected proteins from 45 organisms for examination (listed in Tab. S1). Of these organisms, 16 are classified as 'Model organisms' within the AFDB. The table presents each organism's species name, Proteome ID, total protein chain length, and number of proteins (with parenthetical notation indicating the subset of proteins with average pLDDT scores exceeding 70).

#### B. Code Availability

The complete codebase for all analyses in this study is available on GitHub. It includes implementations for computing structural and dynamical properties of proteins—such as contact order, fluctuation entropy, modularity, fractal dimension, and renormalization procedures—along with step-by-step documentation and usage examples to support replication. We also share all computed results of the structure- and dynamics-related measures to ensure full reproducibility of our findings.

TABLE S1. Overview of the protein structures used in this study, all sourced from AlphaFold Database (AFDB).

| Organism | Proteome ID | Total Length | # Protein | Label |
|---|---|---|---|---|
| *Helicobacter pylori* | UP000000429_85962 | 491886 | 1554(1425) | |
| *Methanocaldococcus jannaschii* | UP000000805_243232 | 505519 | 1787(1684) | Model |
| *Campylobacter jejuni* | UP000000799_192222 | 507610 | 1623(1583) | |
| *Haemophilus influenzae* | UP000000579_71421 | 527721 | 1704(1624) | |
| *Mycobacterium leprae* | UP000000806_272631 | 538637 | 1603(1423) | |
| *Neisseria gonorrhoeae* | UP000000535_242231 | 571581 | 2106(1827) | |
| *Streptococcus pneumoniae* | UP000000586_171101 | 587038 | 2030(1943) | |
| *Staphylococcus aureus* | UP000008816_93061 | 797397 | 2889(2694) | |
| *Shigella dysenteriae* | UP000002716_300267 | 1094416 | 3897(3656) | |
| *Mycobacterium tuberculosis* | UP000001584_83332 | 1332031 | 3995(3662) | |
| *Escherichia coli* | UP000000625_83333 | 1354446 | 4403(4215) | Model |
| *Salmonella typhimurium* | UP000001014_99287 | 1421481 | 4533(4340) | |
| *Klebsiella pneumoniae* | UP000007841_1125630 | 1646261 | 5728(5348) | |
| *Pseudomonas aeruginosa* | UP000002438_208964 | 1857268 | 5563(5392) | |
| *Schizosaccharomyces pombe* | UP000002485_284812 | 2379146 | 5117(3938) | Model |
| *Nocardia brasiliensis* | UP000006304_1133849 | 2750577 | 8414(7951) | |
| *Saccharomyces cerevisiae* | UP000002311_559292 | 2937414 | 6060(4264) | Model |
| *Candida albicans* | UP000000559_237561 | 2981594 | 6035(4377) | Model |
| *Paracoccidioides lutzii* | UP000002059_502779 | 3951676 | 8811(4947) | |
| *Dracunculus medinensis* | UP000274756_318479 | 4070558 | 10868(7190) | |
| *Plasmodium falciparum* | UP000001450_36329 | 4095522 | 5361(2449) | |
| *Ajellomyces capsulatus* | UP000001631_447093 | 4155681 | 9214(5092) | |
| *Wuchereria bancrofti* | UP000270924_6293 | 4298790 | 13000(7609) | |
| *Trypanosoma brucei* | UP000008524_185431 | 4332330 | 8561(5112) | |
| *Sporothrix schenckii* | UP000018087_1391915 | 4783716 | 8673(5738) | |
| *Madurella mycetomatis* | UP000078237_100816 | 4853850 | 9733(6957) | |
| *Onchocerca volvulus* | UP000024404_6282 | 4984676 | 12111(6829) | |
| *Leishmania infantum* | UP000008153_5671 | 5116451 | 8045(4479) | |
| *Cladophialophora carrionii* | UP000094526_86049 | 5263867 | 11181(7500) | |
| *Brugia malayi* | UP000006672_6279 | 5473733 | 11551(5485) | |
| *Schistosoma mansoni* | UP000008854_6183 | 5827960 | 9652(7323) | |
| *Strongyloides stercoralis* | UP000035681_6248 | 5851717 | 12864(8316) | |
| *Fonsecaea pedrosoi* | UP000053029_1442368 | 6141607 | 12525(9278) | |
| *Dictyostelium discoideum* | UP000002195_44689 | 6843544 | 12718(7272) | Model |
| *Drosophila melanogaster* | UP000000803_7227 | 7402101 | 13824(8878) | Model |
| *Caenorhabditis elegans* | UP000001940_6239 | 8134629 | 19827(13694) | Model |
| *Trypanosoma cruzi* | UP000002296_353153 | 9670826 | 19242(10546) | |
| *Arabidopsis thaliana* | UP000006548_3702 | 11116446 | 27448(18990) | Model |
| *Homo sapiens* | UP000005640_9606 | 11415704 | 20597(15364) | Model |
| *Mus musculus* | UP000000589_10090 | 11750371 | 21701(15450) | Model |
| *Rattus norvegicus* | UP000002494_10116 | 11762922 | 22897(15111) | Model |
| *Oryza sativa* | UP000059680_39947 | 13413314 | 43667(20928) | Model |
| *Zea mays* | UP000007305_4577 | 14541968 | 39228(21869) | Model |
| *Danio rerio* | UP000000437_7955 | 15032210 | 25777(17995) | Model |
| *Glycine max* | UP000008827_3847 | 21730487 | 55855(35950) | Model |

### II. VALIDITY OF ALPHAFOLD-PREDICTED STRUCTURES FOR STATISTICAL ANALYSIS

Our framework examines topology–dynamics relationships in proteins, linking residue–residue contact topology to both folding dynamics and native-state slow modes. Because these large-scale dynamical properties are governed mainly by the global contact network rather than precise atomic coordinates, our analyses are robust to small local structural inaccuracies. This contact-based perspective offers a consistent basis for statistical comparisons across large datasets, while acknowledging that fine-grained details—such as side-chain conformations or solvent effects—are not explicitly resolved.

To verify that AlphaFold (AF) predictions remain suitable as the structural basis for topology-driven analyses, we examined proteins from the Protein Structural Change DataBase (PSCDB) with experimentally determined apo and holo structures. For each protein, we prepared three structures: (i) the experimental apo form, (ii) the experimental holo form, and (iii) the AF2-predicted structure from the amino acid sequence. We then computed the displacement vectors $\Delta\vec{r}_{\text{AF-apo}}$, $\Delta\vec{r}_{\text{AF-holo}}$, and $\Delta\vec{r}_{\text{holo-apo}}$ (Fig. S1 inset), and evaluated their pairwise cosine similarities.

Across PSCDB, the average maximum cosine similarity was $\sim 0.75$ (Fig. S1), far above the near-zero value expected for random vectors in high-dimensional space. This consistently high alignment indicates that AF-predicted structures often lie close to the natural apo–holo transition pathway, despite potential local inaccuracies. Consequently, the associated NMA-derived slow-mode dynamics remain robust. More broadly, AF-predicted structures frequently reside along functional transition pathways, as also noted in Ref. [1], thereby encoding dynamically relevant conformations and serving as reliable reference points for studying protein motions.

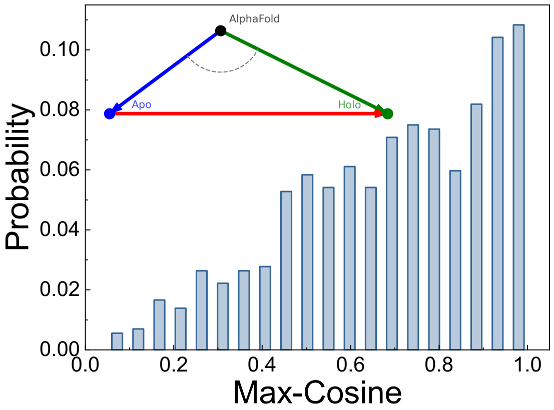


FIG. S1. Cosine similarity between AF-predicted and experimental displacement vectors for proteins with known apo–holo transitions. The mean similarity of $\sim 0.75$ indicates that AF predictions often lie close to natural transition pathways.

### III. STRUCTURE- AND SEQUENCE-BASED ENTROPY MEASURES

We use two complementary entropy measures to jointly characterize flexibility and evolutionary variability: fluctuation entropy $S$ from the AF2-derived contact network, and sequence entropy $H$ obtained independently from the nonlinear amino-acid probability distributions of protein language models, such that any agreement between them does not arise from shared modeling assumptions. In this section, we describe the model setups and computation procedures for both entropy measures.

#### A. Fluctuation Entropy from Gaussian Network Model (GNM)

The Gaussian Network Model (GNM) is a coarse-grained approach for analyzing protein dynamics where amino acid residues are represented as nodes, typically located at their $C_\alpha$ atom positions. This simplified representation reduces the computational complexity while retaining the essential physical properties governing protein dynamics. In GNM, pairs of nodes within a cutoff distance $r_c$ are connected by harmonic springs with uniform force constants, creating a network that captures the topology of inter-residue contacts in the native structure. The model is defined by the Kirchhoff (or Laplacian) matrix $L$ of the residue contact network:

$$L_{ij} = \begin{cases} -1 & \text{if } r_{ij} < r_c \text{ and } i \neq j \\ -\sum_{k\neq i} L_{ik} & \text{if } i = j \\ 0 & \text{otherwise} \end{cases} \tag{S1}$$

The diagonal entry $L_{ii}$ equals the degree $\deg(i)$ of node $i$, representing the total number of connections involving residue $i$. For simplicity, we set the cutoff distance $r_c$ for the GNM equal to that used in the calculation of contact order, i.e., $r_c = r_{\text{CO}}$, so that the same geometric criterion defines both contact edges in the network and the set of total contacts. However, it is worth noting that elastic network models such as the GNM include both short-range ($|i-j| < 4$) and long-range ($|i-j| \geq 4$) contacts, thereby encompassing local backbone conductivities and tertiary-level residue contacts based on spatial proximity.

The key relationship in GNM is that the covariance matrix $C$ of residue fluctuations is proportional to the pseudoinverse of the Laplacian matrix [2, 3]:

$$C = k_B T \cdot L^+, \tag{S2}$$

where $k_BT$ is the thermal energy and $L^+$ is the pseudoinverse of $L$. For a connected graph—valid for proteins with continuous structural connectivity—the Laplacian $L$ has a zero eigenvalue $\sigma_0 = 0$ corresponding to the eigenvector $\mathbf{1}$ (a column vector of all ones). This zero mode arises because each row of $L$ sums to zero and represents global translation. The remaining nonzero eigenvalues are ordered as $0 < \sigma_1 \leq \sigma_2 \leq \cdots \leq \sigma_{N-1}$. The covariance matrix $C$ has eigenvalues in the reverse order, $\lambda_1 \geq \lambda_2 \geq \cdots \geq \lambda_{N-1}$, and a corresponding zero eigenvalue $\lambda_0 = 0$. The pseudoinverse $L^+$ is constructed from the nonzero eigenvalues $\sigma_k$ and their eigenvectors $v_k$:

$$L^+ = \sum_{k=1}^{N-1} \frac{1}{\sigma_k} v_k v_k^T. \tag{S3}$$

The covariance eigenvalues relate to the Laplacian via $\lambda_k = k_BT/\sigma_k$. The fluctuation entropy, defined as $S = \log\det C = \sum_{k=1}^{N-1} \log\lambda_k$, simplifies to $S = -\sum_{k=1}^{N-1} \log\sigma_k$ when $k_BT = 1$, directly linking network topology to conformational flexibility [4–6]. The derivation is given below:

In the harmonic approximation, the conformational distribution around the native state follows a multivariate Gaussian:

$$P(\Delta\vec{r}) = \frac{1}{Z} \exp\left(-\frac{1}{2k_BT} \Delta\vec{r}^T L \Delta\vec{r}\right), \tag{S4}$$

where $Z$ is the normalization constant, and $\Delta\vec{r} = [\Delta\vec{r}_1, \Delta\vec{r}_2, \cdots \Delta\vec{r}_N]$ is a high dimensional vector describing the deformations of residues. The Shannon entropy of this distribution in the standard form is:

$$S = -\int P(\Delta\vec{r}) \ln P(\Delta\vec{r}) d(\Delta\vec{r}) \tag{S5}$$

For a multivariate Gaussian distribution, this entropy evaluates to:

$$S = \frac{1}{2} \ln \left( (2\pi e)^{N-1} \det(C) \right), \tag{S6}$$

where $C$ is the covariance matrix, and det denotes the pseudodeterminant, i.e., the product of its nonzero eigenvalues. Since $C \propto L^{+}$ and $L$ has exactly one zero eigenvalue, we get:

$$S = \frac{1}{2} \ln \left( (2\pi e)^{N-1} (k_B T)^{N-1} \prod_{k=1}^{N-1} \frac{1}{\sigma_k} \right). \tag{S7}$$

Simplifying and dropping constant terms:

$$S \propto -\sum_{k=1}^{N-1} \ln \sigma_k = \sum_{k=1}^{N-1} \log \lambda_k, \tag{S8}$$

which quantifies the logarithmic volume of accessible conformational space.

In this study, we focus on the $k = 8$ lowest-frequency modes (those with the largest $\lambda_k$), which dominate conformational variance and capture the most collective, functionally relevant motions:

$$S_k \propto \sum_{t=1}^{k} \log \lambda_t. \tag{S9}$$

Varying $k$ does not affect the observed CO–$S$ scaling. More generally, fluctuation entropy can be computed over any selected subset of modes $\mathcal{M}$,

$$S_{\mathcal{M}} \propto \sum_{t \in \mathcal{M}} \log \lambda_t, \tag{S10}$$

allowing separate analysis of global domain motions, localized fluctuations, or structurally constrained regions, and thereby isolating the contributions of different dynamical components to the total entropy.

#### B. Sequence Entropy from Pretrained Protein Language Model

As mentioned in the main text, sequence entropy $H$ is computed using the pretrained protein language model `esm2_t30_150M_UR50D`, which estimates site-specific amino acid distributions from sequence context alone, without using structural information. For a protein of length $N$, the model provides, at each position $i$,

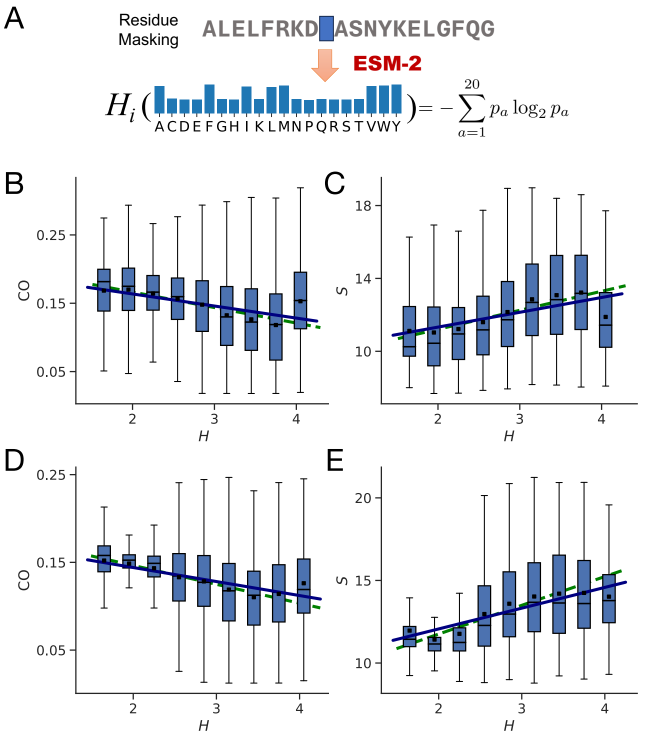


FIG. S2. (A) Schematic of sequence entropy computation using ESM2. (B–C) CO–$H$ and $S$–$H$ relationships for proteins with $175 \le N < 225$. Box plots are grouped by sequence entropy $H$ (outliers omitted). For both quantities, binned means (dots with solid-line fits) and fits to the full data (dashed lines) closely coincide, indicating consistent negative CO–$H$ and positive $S$–$H$ correlations. (D–E) Same analysis for proteins with $275 \le N < 325$, showing the same robust trends across size ranges.

the conditional distribution $p_i(a \mid \text{context})$ over the 20 amino acids. The site-wise entropy is

$$H_i = -\sum_{a \in \mathcal{A}_{20}} p_i(a \mid \text{context}) \log_2 p_i(a \mid \text{context}), \tag{S11}$$

and the sequence entropy is the positional average $H = \frac{1}{N} \sum_{i=1}^{N} H_i$. Here, with values ranging from 0 (fully conserved; one amino acid strongly preferred) to $\log_2 20 \approx 4.3$ bits (all amino acids equally likely). Thus, larger $H_i$ indicates positions that tolerate more substitutions, while smaller values reflect stronger evolutionary constraints, providing a sequence-based measure of mutational variability independent of any structural model.

In the main text, we reported the CO–$H$ and $S$–$H$ relationships for proteins with $225 \le N < 275$. Here, we additionally show the same relationships for proteins with $175 \le N < 225$ (panels B–C) and $275 \le N < 325$ (panels D–E), confirming that the observed trends are robust across size ranges. A slight deviation from the overall trend is observed in the highest-$H$ bin in the figures. This arises from the use of AlphaFold-predicted structures subject to a pLDDT-based confidence filter, which preferentially excludes large proteins with extremely high sequence entropy. Including these excluded sequences restores consistency with the overall trend, indicating that the deviation reflects a sampling effect rather than a breakdown of the underlying relationship. Notably, $H$ is derived entirely from residue probability distributions learned from large-scale evolutionary sequence data using a nonlinear protein language model. In this sense, $H$ provides an evolutionary counterpart to the structure-derived fluctuation entropy $S$.

#### C. Robustness of Contact Order and Fluctuation Entropy to Parameter Variations

In the GNM analysis, we use a cutoff distance of $r_c = r_{\mathrm{CO}} = 8$ Å for calculating both contact order (CO) and fluctuation entropy $S$. To simplify computation, $S$ was primarily estimated using the first 8 nonzero modes. Here, we demonstrate that our results are robust with respect to both the choice of cutoff distance and the number of modes considered in the calculation of $S$. Similar to the Main Text, we conduct the analysis for the proteins with similar chain lengths ($225 \leq N < 275$) and mean pLDDT $\geq 70$. As shown in Fig. S3A–C, varying the contact cutoff distance used to compute CO ($r_{\mathrm{CO}} = 7, 9$, and 10 Å) results in values that are linearly proportional to those obtained with the standard $r_{\mathrm{CO}} = 8$ Å. Similarly, Fig. S3D–F shows that when varying the cutoff distance used to construct the GNM network ($r_c = 7, 9$, and 10 Å), the resulting $S$ remains proportional to that calculated with $r_c = 8$ Å. These results support the robustness of our Main Text analysis with respect to parameter choices and the number of modes included in estimating $S$.

Using the same group of proteins, we further validated that the number of modes included in the calculation does not qualitatively affect the results. Specifically, we computed fluctuation entropy $S$ using a fixed cutoff distance $r_c = 8$ Å, while varying the number of modes considered. As shown in Fig. S4A–E, we compared results based on the contributions from the first 1, 2, 4, 16, and all $N - 1$ nonzero modes. Across all comparisons, the resulting $S$ values remain linearly proportional, indicating that while the absolute magnitude of $S$ depends on the number of modes included, the relative ordering and correlations remain consistent. This confirms the robustness of our Main Text results with respect to mode selection in the entropy calculation.

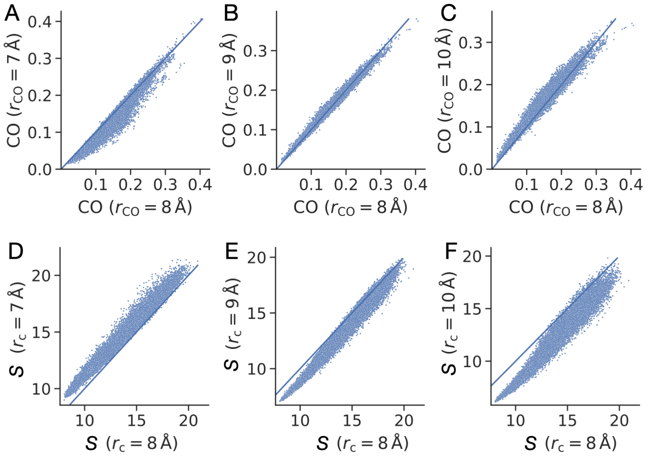


FIG. S3. Robustness of CO and $S$ under different parameter choices. (A–C) CO values computed with contact cutoffs $r_{\mathrm{CO}} = 7, 9$, and 10 Å are linearly proportional to those obtained with the standard $r_{\mathrm{CO}} = 8$ Å. (D–F) Fluctuation entropy $S$, computed using the first 8 modes and GNM cutoffs $r_c = 7, 9$, and 10 Å, remains linearly proportional to the reference case with $r_c = 8$ Å. In all panels, the blue line denotes the identity line $x = y$, highlighting consistency across parameter settings.

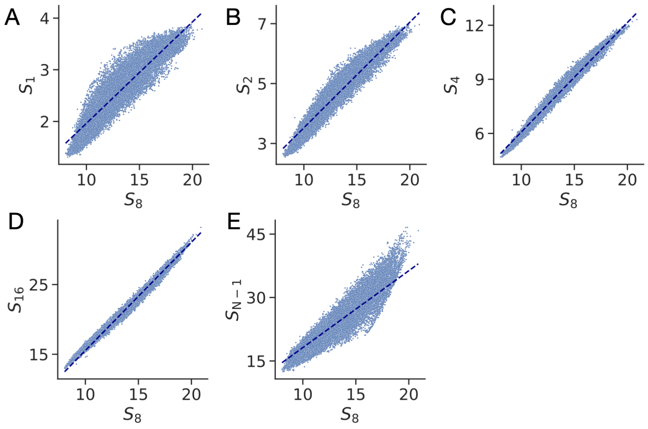


FIG. S4. Robustness of fluctuation entropy $S$ to the number of modes included in the GNM calculation. Using a fixed cutoff distance $r_c = 8$ Å, $S_k$ denotes the entropy computed using the first $k = 1, 2, 4, 16$, and $N-1$ nonzero modes, respectively (A–E). Each panel shows the relationship between $S_k$ and the reference $S$ (i.e., $S_8$, as used in the Main Text). The blue dash line indicates a linear fit between $S_k$ and $S_8$, demonstrating strong proportionality. These results confirm that although the absolute magnitude of entropy varies with the number of modes, the relative trends are preserved.

#### D. Consistency between Normal Mode Analysis and AlphaFold uncertainty measures

Normal mode analysis (NMA) provides a topology-based, harmonic approximation of protein dynamics, enabling consistent fluctuation entropy ($S$) estimates across large datasets. To evaluate its validity, we compared NMA-derived fluctuations with AF prediction confidence metrics—pLDDT and Predicted Aligned Error (PAE).

For >800 proteins from the Protein Structural Change DataBase (PSCDB), NMA fluctuations from AF-predicted structures showed strong correlations (Pearson/Spearman 0.5–0.7) with $100 - $ pLDDT (Fig. S5). Peaks in NMA fluctuations matched regions of low AF confidence, robust to changes in cutoff distance and number of modes. Across diverse proteins, $S$ also correlated strongly with total PAE (sum over the full matrix; Fig. S6), indicating that $S$ reliably captures global flexibility consistent with AF uncertainty predictions.

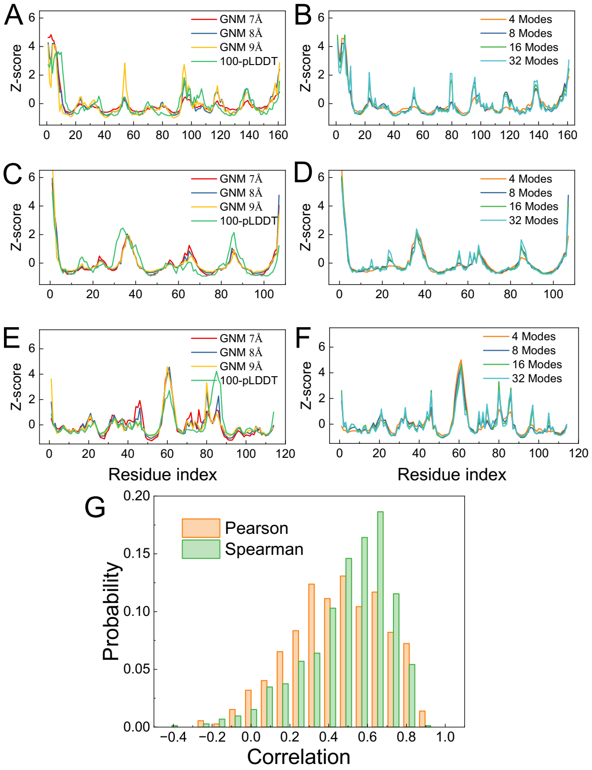


FIG. S5. Comparison between NMA-predicted residue fluctuations and $100 - \text{pLDDT}$ for GroES (A,B), HRAS (C,D), and HSP70 (E,F), normalized as $z$-scores. (G) Distribution of Pearson and Spearman correlations for PSCDB proteins.

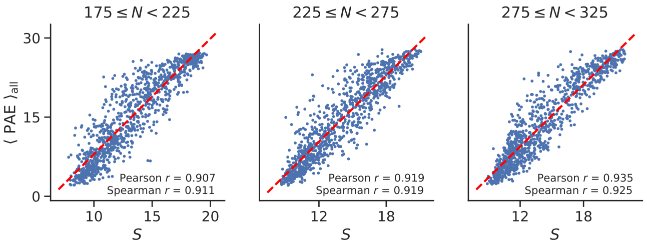


FIG. S6. Correlation between fluctuation entropy $S$ and total PAE from AF-predicted structures across proteins of varying chain lengths $N$.

#### E. Extension to Anisotropic Network Model (ANM)

The GNM can be generalized to the Anisotropic Network Model (ANM), which incorporates directional information about residue fluctuations [7]. In ANM, the Hessian matrix $H$ is a $3N \times 3N$ matrix describing the second derivatives of the potential energy with respect to Cartesian coordinates. Similar to GNM, there exists an inverse relationship between the covariance matrix and the Hessian: $C = k_B T \cdot H^+$. The Hessian has six zero eigenvalues corresponding to rigid-body motions (three translations and three rotations). The fluctuation entropy calculation follows the same principle as in GNM but excludes these six zero modes when computing the pseudodeterminant.

### IV. STRUCTURAL BASIS OF THE CO–$S$ RELATIONSHIP WITH DYNAMICAL AND FUNCTIONAL IMPLICATIONS

#### A. Representative Protein Examples Illustrating the CO–$S$ Relationship

| | UNIPROT | Species Code | $N$ | $R_g$/Å | CO | $S$ |
|---|---|---|---|---|---|---|
| (A) | A0A0D2F035 | 9EURO2 | 264 | 18.07 | 0.283 | 9.10 |
| (B) | Q0PBA2 | CAMJE | 260 | 24.71 | 0.124 | 12.07 |
| (C) | A0A044V453 | ONCVO | 263 | 43.12 | 0.049 | 15.32 |
| (D) | P34067 | RAT | 263 | 22.82 | 0.194 | 13.80 |
| (E) | Q8ZRP4 | SALTY | 274 | 20.54 | 0.088 | 10.64 |

TABLE S2. Selected proteins representing different combinations of contact order (CO) and fluctuation entropy ($S$). All proteins have similar lengths ($N \approx 260$), allowing a direct comparison of structural topology ($CO$), compactness (radius of gyration $R_g$), and native-state dynamics ($S$). The calculations are based on cutoff distances $r_c = r_{\text{CO}} = 8$ Å, and the fluctuation entropy $S$ includes contributions from the first eight leading normal modes.

To illustrate the relationship between protein topology and dynamics, we present five representative proteins spanning a range of contact order (CO) and fluctuation entropy $S$ values (Table S2, Figure S7). These examples were selected to highlight distinct structural regimes: for instance, protein A0A0D2F035 (panel A) has high CO and low $S$, consistent with a compact fold stabilized by dense long-range contacts, while A0A044V453 (panel C) exhibits very low CO and high $S$, reflecting an extended structure with high conformational flexibility. Proteins such as Q8ZRP4 (panel E), with both low CO and low $S$, likely represent short-range-dominated folds with more rigid native states. All five proteins have similar sequence lengths ($N \approx 260$), enabling a direct comparison of their structural and dynamic characteristics.

#### B. Contact density as a source of scatter in the CO–$S$ relation

Although CO and $S$ follow a clear global trend, systematic deviations from this relation are largely explained by

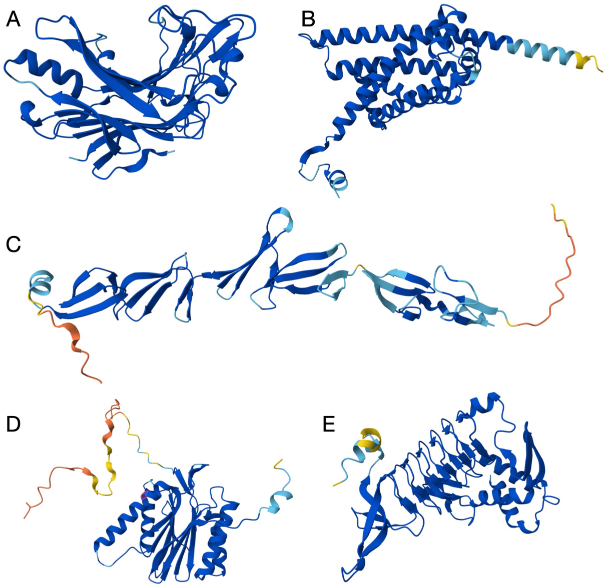


FIG. S7. Cartoon representations of the five proteins listed in Table S2, with backbone coloring based on AF2 pLDDT scores (blue: high confidence; red/orange: low confidence). The proteins are in the same order as listed in the table.

differences in the total number of residue–residue contacts $N_c$. To examine this effect, we selected proteins of similar chain length ($225 \leq N < 275$) and grouped them into three narrow contact-order bins: CO $\approx 0.10$ ($0.0925 \leq$ CO $< 0.1075$), CO $\approx 0.15$ ($0.1425 \leq$ CO $< 0.1575$), and CO $\approx 0.20$ ($0.1925 \leq$ CO $< 0.2075$). Within each bin, CO values are nearly identical, so variations in $S$ primarily reflect other structural features.

As shown in Figure S8, $S$ decreases systematically with increasing $N_c$ in all three groups. Proteins with fewer contacts, corresponding to higher native-state energy, exhibit greater fluctuation entropy due to weaker topological constraints and access to a broader conformational ensemble. This trend is consistent with the funneled energy landscape picture, in which contact density regulates the accessible conformational space.

### C. Anharmonic protein motions constrained by long-range contacts

To validate, using independent structural datasets, that large-scale anharmonic protein motions are predominantly governed by a specific subset of residue–residue contacts, we performed the following analysis. Based on the proposed CO–$S$ relation, we hypothesize that contacts with large sequence separations impose the strongest constraints on global conformational flexibility, and that their selective removal disproportionately amplifies slow-mode amplitudes beyond harmonic predictions.

We first tested this using 1,000 AF-predicted proteins of similar chain length ($N \approx 250$). As shown in Fig. S9A, removing short-range contacts produced only minor increases in $S$, whereas removing long-range contacts caused substantial increases, indicating that long-range interactions are the primary constraints on large-scale, potentially anharmonic conformational changes.

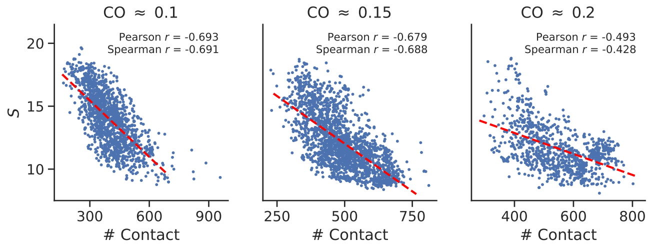


FIG. S8. For proteins of similar chain length ($225 \leq N < 275$), fluctuation entropy $S$ versus number of contacts $N_c$ in three narrow contact-order bins: CO $\approx 0.10$, CO $\approx 0.15$, and CO $\approx 0.20$. Bin widths are chosen so that CO values are effectively constant within each group. In all cases, $S$ decreases with $N_c$, indicating that deviations from the global CO–$S$ trend in Figure 2A are largely attributable to variations in contact density.

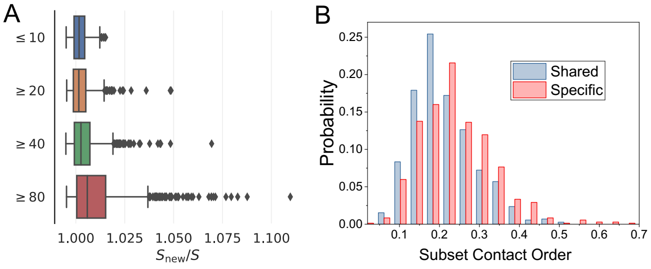


FIG. S9. (A) Change in fluctuation entropy $S$ after removing 5% of contacts grouped by sequence separation in 1,000 AF-predicted proteins of similar chain length ($N \approx 250$). Short-range contacts correspond to separation $\leq 10$ residues; long-range contacts have separation $\geq 20$, $\geq 40$, or $\geq 80$ residues. Removing short-range contacts causes only minor increases in $S$, whereas removing long-range contacts produces substantial increases, indicating their dominant role in constraining large-scale motions. (B) For PSCDB proteins with experimentally observed apo–holo transitions, shared contacts occur mainly at low Sub-CO, while state-specific contacts occur at higher Sub-CO and are preferentially broken or formed during transitions, highlighting the disproportionate influence of long-range constraints on functional rearrangements.

We further validated this relationship using proteins from PSCDB [8] with experimentally observed functional transitions. Here, we introduced the *Subset Contact Order* (Sub-CO)—the average sequence separation of a defined contact subset, normalized by chain length $N$—as a structural descriptor linking contact topology to motion amplitude. Following Li *et al.* [9], contacts were classified as *shared* (present in both apo and holo) or *state-specific* (present in only one state), and their Sub-CO values compared (Fig. S9B). Shared contacts predominantly exhibit low Sub-CO, consistent with maintaining local geometry in both states, whereas state-specific contacts have higher Sub-CO and are preferentially broken or formed during transitions. This pattern mirrors our selective-removal results and agrees with previous studies [10–12] that emphasize the dominant role of long-range constraints in large-scale rearrangements, highlighting the limitations of purely harmonic models and the need to incorporate anharmonic effects.

Overall, the results confirm our hypothesis: long-range constraints—captured by high Sub-CO—act as structural bottlenecks for large-scale, anharmonic motions, and their targeted disruption drives functional rearrangements by selectively releasing topological constraints. High Sub-CO contacts thus emerge as both a structural hallmark and a functional driver of large-amplitude conformational changes. Their removal not only directly increases entropy but may also trigger a cascade-like loss of other long-range constraints, further amplifying structural flexibility. Incorporating such cooperative effects into anharmonic models represents an important future direction for understanding and predicting protein functional dynamics.

#### D. Correlation Between the CO–$S$ Relationship and Biological Function

In this subsection, we examine how the CO–$S$ relationship varies with biological function, using the Gene Ontology (GO) framework [13, 14], which classifies proteins into standardized functional categories. Our analysis employs the simplified GO Slim subset, which provides a reduced vocabulary covering three main aspects: *Biological Process* (e.g., cell cycle, metabolic process), *Cellular Component* (e.g., plasma membrane, extracellular), and *Molecular Function* (e.g., binding, catalytic activity). Although GO Slim omits mechanistic details, it enables tractable, large-scale statistical comparisons. A single protein may belong to multiple GO Slim categories due to the presence of multiple functional domains, and GO terms are hierarchical. Here, we focus on one level below the top-level categories to preserve meaningful functional resolution while keeping the analysis general. To minimize chain-length effects, we restrict the dataset to model-organism proteins with $225 \leq N < 275$, ensuring that CO and $S$ values are directly comparable across groups.

We compared the distributions of CO and $S$ across GO Slim categories using box plots and Kolmogorov–Smirnov (KS) tests. The distributional comparisons are shown in Fig. S10 (panel A: CO; panel B: $S$), while the full statistical significance map is presented in Fig. S11 (panel A: CO; panel B: $S$), where asterisks denote $p$-values and color encodes KS statistics. The most notable deviations of functional subgroups from their broader categories reveal distinct topological–dynamical profiles:

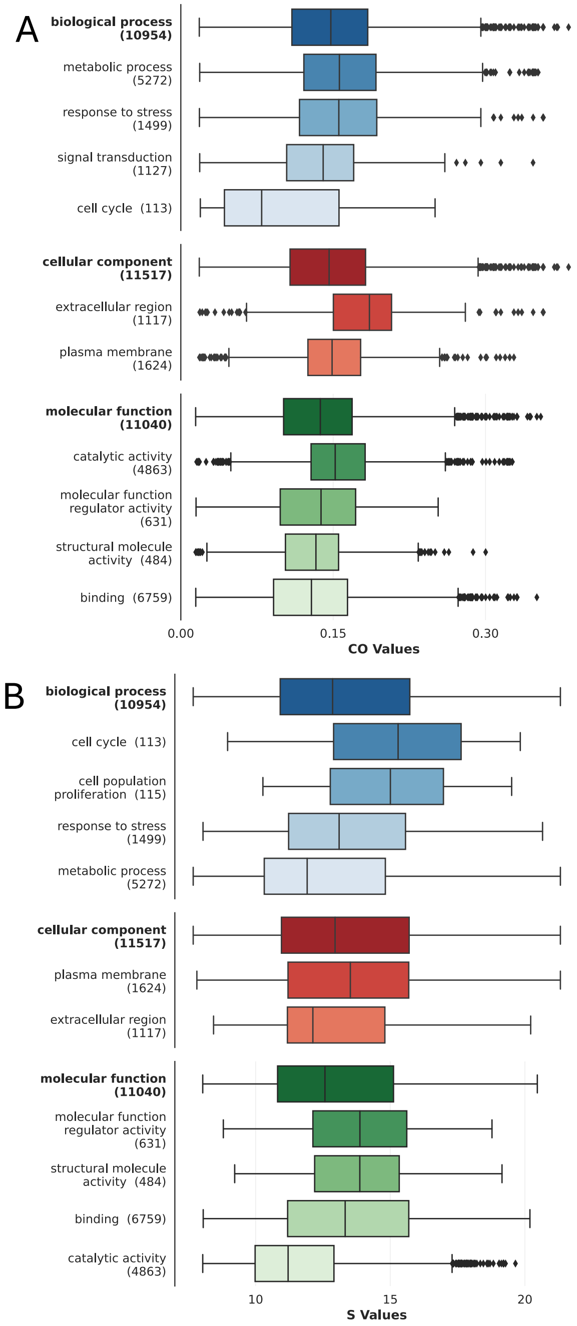


FIG. S10. Distributions of contact order (CO, panel A) and fluctuation entropy ($S$, panel B) across GO Slim categories for model-organism proteins with $225 \leq N < 275$. Distinct functional categories exhibit characteristic topological–dynamical profiles.

**(1) Biological Process.** Proteins annotated under "cell cycle" tend to exhibit significantly lower CO and higher $S$, reflecting the need for substantial conformational flexibility to support regulated structural rearrangements during cell-cycle progression. By contrast, proteins associated with "metabolic process" generally display higher CO and lower $S$, indicating more rigid architectures that

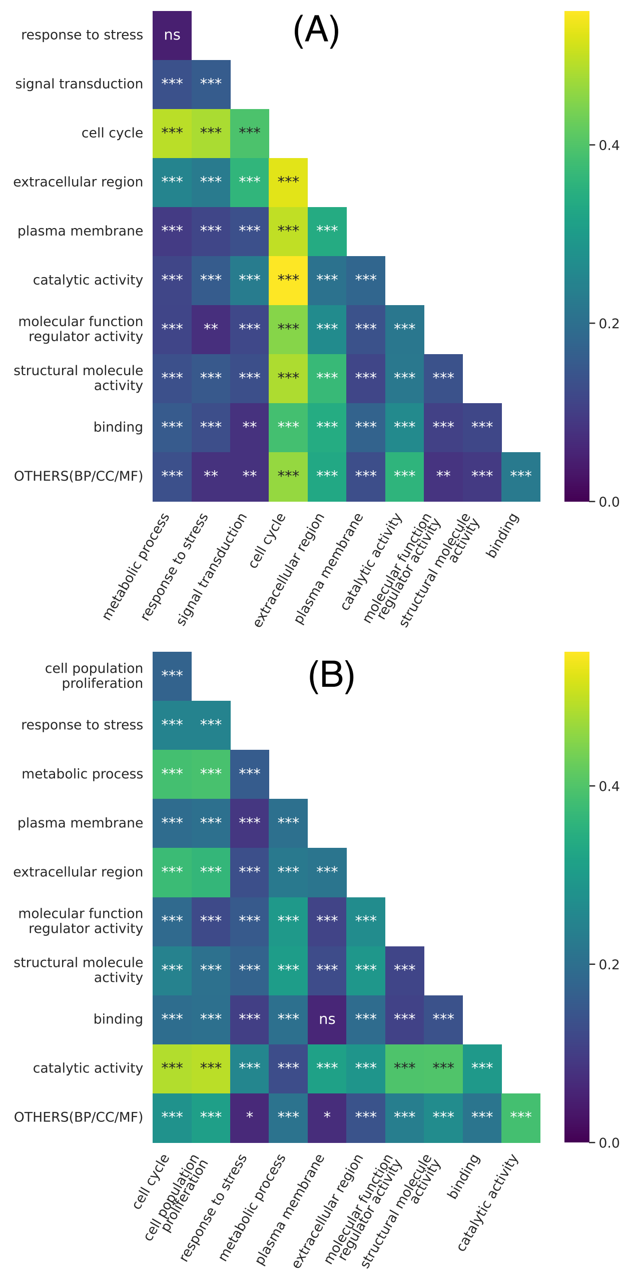


FIG. S11. Pairwise statistical comparisons of CO (panel A) and $S$ (panel B) across GO Slim categories. Color encodes KS statistics; asterisks indicate significance levels based on $p$-values.

help maintain well-defined conformations suited for catalytic or otherwise structurally stable functions.

**(2) Molecular Function.** Proteins classified under "catalytic activity" typically show higher CO and lower $S$ compared to other molecular function categories. This suggests that high-contact-order topologies contribute to stabilizing active-site geometry and restricting conformational fluctuations. Here, "catalytic activity" is used in a broad sense, encompassing proteins that facilitate chemical transformations, not only canonical enzymes.

Other categories may also show significant deviations from their respective baselines, indicating that functional specialization is associated with characteristic topological–dynamical profiles. These results support the idea that the CO–$S$ trend identified in the main text is modulated by evolutionary adaptation to functional requirements. While exploratory, this analysis demonstrates the potential for linking topological–dynamical metrics to functional annotation, offering a pathway to connect structural physics with biological roles.

### V. A GRAPH-THEORETICAL PERSPECTIVE ON THE CO–$S$ RELATIONSHIP

#### A. Intuitive Picture

Before proceeding to the formal derivation, we aim to establish a general physical understanding of the connection between fluctuation entropy and spanning tree number. A spanning tree is a subset of edges that connects all nodes without forming any cycles. To illustrate this concept, consider the examples shown in Fig. S12. Panel A presents a small network with five nodes and five edges containing one cycle. In this configuration, there exist exactly five possible spanning trees. This is because removing any single edge from the cycle transforms the structure into a tree. All possible spanning tree configurations for this example are illustrated in Panel B. Panel C depicts another configuration with the same number of nodes and edges, but with a different topology. Here, nodes 2, 3, and 4 form a smaller cycle. To convert this graph into a tree, we have only three options—removing any edge from this smaller cycle. The corresponding spanning tree configurations are shown in Panel D. These examples demonstrate that, given a similar number of edges, networks with fewer and smaller cycles yield a reduced number of possible spanning trees. This intuitive relationship aligns with the principles illustrated in Fig. 2B and 2C of the Main Text.

#### B. Formal Proof

In this subsection, we formalize the relationship between the topological structure of a protein's contact network and its fluctuation entropy. We begin by relating entropy to the number of spanning trees via the Laplacian spectrum. Then, we show how the placement of a fixed number of non-local contacts affects the effective resistance between residues and the resulting fluctuation entropy. Finally, we establish that higher contact order leads to lower entropy by biasing edge placement toward residue pairs with higher resistance distance.

Throughout this analysis, we fix the number of residues $N$, and compare structures with equal numbers of contacts (edges). Additive constants such as $\log N$ in the

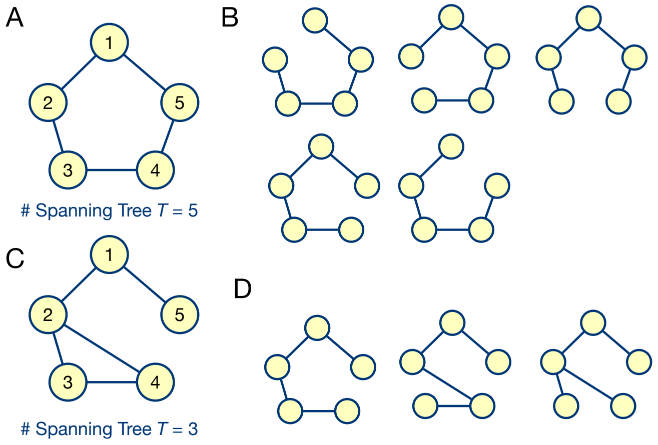


FIG. S12. Spanning trees in network configurations. (A) A 5-node network with one cycle, yielding five possible spanning trees (B) by removing one edge from the cycle. (C) A 5-node network with a smaller 3-node cycle, yielding only three spanning trees (D). These examples demonstrate how networks with fewer or smaller cycles generate fewer spanning trees, paralleling the relationship between structural rigidity and fluctuation entropy (Main Text, Fig. 2B,C).

Matrix-Tree Theorem are absorbed into a baseline and omitted in comparative entropy statements.

**Theorem 1.** *For a protein structure modeled as an elastic network with $N$ nodes, the fluctuation entropy $S$ and the number of spanning trees $T$ satisfy:*

$$S = -\log T + \textit{const}. \tag{S12}$$

*Proof.* Let $L$ be the Laplacian matrix of the elastic network, and let $\{\sigma_k\}_{k=1}^{N-1}$ be its non-zero eigenvalues. The fluctuation entropy is given by:

$$S = -\sum_{k=1}^{N-1} \log \sigma_k.$$

By the Matrix-Tree Theorem [15–17], the number of spanning trees $T$ is:

$$T = \frac{1}{N} \prod_{k=1}^{N-1} \sigma_k.$$

Taking logarithms:

$$\log T = \sum_{k=1}^{N-1} \log \sigma_k - \log N \quad \Rightarrow \quad S = -\log T - \log N.$$

For fixed $N$, the term $\log N$ is constant, so we absorb it into a baseline:

$$S = -\log T + \text{const}.$$

□

**Theorem 2** (Edge Placement Affects Entropy Under Fixed Edge Count)**.** *Given two elastic networks with the same number of nodes and the same total number of edges, placing a contact between residues with higher effective resistance increases the number of spanning trees and hence the fluctuation entropy.*

*Proof.* Let $G$ be a base graph (e.g., a locally connected backbone), and suppose we are allowed to add one extra edge to it while keeping the total number of edges fixed (e.g., by removing another edge).

Let $G_1 = G + (i,j)$ and $G_2 = G + (k,\ell)$ be two such graphs, each with the same number of edges, where $(i,j)$ is a non-local contact and $(k,\ell)$ is a local contact. Then the number of spanning trees in each graph satisfies:

$$\frac{T_{G_1}}{T_{G_2}} \approx \frac{1 + R_{\text{eff}}(i,j)}{1 + R_{\text{eff}}(k,\ell)}.$$

Here, $R_{\text{eff}}(i,j)$ is the effective resistance between nodes $i$ and $j$, defined as:

$$R_{\text{eff}}(i,j) = (e_i - e_j)^\top L^+ (e_i - e_j),$$

where $L^+$ is the Moore–Penrose pseudoinverse of the Laplacian matrix and $e_i$ is the standard basis vector. Physically, $R_{\text{eff}}(i,j)$ quantifies how easily fluctuations, energy, or information propagate between residues $i$ and $j$, and is analogous to the electrical resistance between two nodes in a resistor network where each edge represents a unit resistor.

In a sparse graph like a protein backbone, non-local pairs $(i,j)$ are typically farther apart and have higher $R_{\text{eff}}$ than local pairs $(k,\ell)$, implying:

$$T_{G_1} > T_{G_2} \quad \Rightarrow \quad S_{G_1} < S_{G_2}.$$

Thus, even under a fixed number of edges, placing contacts between high-resistance pairs leads to more spanning trees and higher fluctuation entropy. □

**Theorem 3** (Effective Resistance Grows with Sequence Separation)**.** *In a protein backbone dominated by local contacts, the effective resistance $R_{\text{eff}}(i,j)$ between two residues increases with their sequence separation $|i-j|$.*

*Proof.* To analyze how the effective resistance $R_{\text{eff}}(i,j)$ scales with sequence separation $|i-j|$, we begin with the simplest case: a linear chain (path graph) with unit edge weights. In this case, it is well known that

$$R_{\text{eff}}(i,j) = |i-j|.$$

We now generalize to a broader class of graphs resembling protein backbones—locally connected networks with bounded degree and no long-range shortcuts. In such graphs, each node is connected only to a fixed number of nearby neighbors (e.g., $i\pm1, i\pm2$), and the graph remains sparse. Let $l_G(i,j)$ denote the shortest path length between nodes $i$ and $j$. Since only local contacts are allowed, we have $l_G(i,j) \sim |i-j|$.

From electrical network theory [18, 19], the effective resistance satisfies the lower bound

$$R_{\text{eff}}(i,j) \geq \frac{l_G(i,j)}{\mathcal{D}_{\max}},$$

where $\mathcal{D}_{\max}$ is the maximum degree of the graph. This follows from the Nash–Williams inequality and Rayleigh monotonicity, which imply that in sparse graphs without long-range shortcuts, resistance grows proportionally to distance. Therefore, for large $|i-j|$, we obtain a linear lower bound

$$R_{\text{eff}}(i,j) \geq c|i-j|,$$

for some constant $c > 0$.

On the other hand, since $R_{\text{eff}}(i,j)$ is bounded above by the length of any path from $i$ to $j$, and the number of such paths does not grow exponentially in locally connected structures, resistance cannot fall significantly below the shortest path length. This yields an upper bound of the form

$$R_{\text{eff}}(i,j) \leq C|i-j|,$$

for some constant $C > 0$.

Combining both bounds, we conclude that

$$R_{\text{eff}}(i,j) = \Theta(|i-j|), \quad \text{as } |i-j| \to \infty.$$

Thus, in sparse, locally connected graphs that model protein backbones, the effective resistance between residues grows linearly with their sequence separation in the asymptotic limit. □

**Theorem 4** (Contact Order and Fluctuation Entropy)**.** *Given a protein with contact order* CO *and fluctuation entropy $S$, there exists a negative correlation between* CO *and $S$.*

*Proof.* Let $\mathcal{C}$ denote the set of non-local contacts. For a fixed backbone and total number of contacts, the total number of spanning trees is

$$T = T_0 \prod_{(i,j)\in\mathcal{C}} \left(1 + R_{\text{eff}}(i,j)\right).$$

Taking logarithms gives

$$\log T = \log T_0 + \sum_{(i,j)\in\mathcal{C}} \log(1 + R_{\text{eff}}(i,j)).$$

Since $R_{\text{eff}}(i,j) = \Theta(|i-j|)$, and $\log(1+R_{\text{eff}})$ is monotonic, we can approximate

$$\log T = \log T_0 + \sum_{(i,j)\in\mathcal{C}} f(|i-j|), \quad \text{with } f(\cdot) \text{ increasing.}$$

Thus, higher CO implies greater average $|i-j|$, larger $T$, and therefore lower entropy

$$S = -\log T + \text{const.}$$

□

With the theorems above, we show that for a given chain length $N$, higher CO corresponds to more spanning trees and lower $S$. The derivation is exact when effective resistances are updated sequentially as edges are added, while using backbone resistances provides a good approximation when contacts are sparse and weakly overlapping. Crucially, sparsity is essential and holds for proteins, which can be modeled as geometric graphs, since long-range shortcuts or hubs can collapse resistances and weaken the linear scaling.

#### C. Extension to Weighted Graph Laplacians

The unweighted Laplacian $L = D - A$ treats all contacts equally, but elastic network models with variable interaction strengths are more naturally represented by the weighted Laplacian $L_w$, where edge weights $w_{ij} > 0$ reflect contact strength or geometric proximity. The Matrix-Tree Theorem extends directly to this setting [20], giving the weighted number of spanning trees

$$T_w = \frac{1}{N} \prod_{k=1}^{N-1} \sigma_k^{(w)}, \quad S_w = -\log T_w + \text{const.},$$

where $\{\sigma_k^{(w)}\}$ are the nonzero eigenvalues of $L_w$. The effective resistance generalizes as

$$R_{\text{eff}}^{(w)}(i,j) = (e_i - e_j)^\top L_w^+ (e_i - e_j),$$

preserving its role as a measure of dynamical decoupling between residues. As in the unweighted case, adding contacts between high-resistance pairs increases $T_w$ and reduces entropy. Since high CO biases edge placement toward such distant pairs, the negative CO–$S$ correlation remains valid in the weighted case, confirming the robustness of our conclusions under more realistic protein models.

#### D. Spectral–topological Interpretation of the Critical Behaviors

In this subsection we unify two perspectives—a spectral interpretation based on entropy maximization and a graph-theoretic interpretation grounded in topology—to explain the emergence of scaling laws. Together they show how power-law behavior and underlying contact topology reflect both physical constraints and evolutionary pressures.

Our earlier work [5] demonstrated that the power-law vibrational spectrum arises from a balance between *functional sensitivity* and *mutational robustness*. The former is related to the fluctuation of the protein system and is closely captured by the fluctuation entropy $S$ discussed in this paper. The latter is defined as the robustness of the eigenspace $\mathcal{V}$ spanned by all normal-mode eigenvectors. Perturbation theory shows that the stability of $\mathcal{V}$

is bounded by the minimum eigengap, and is maximized when the spectrum $\{\sigma_i\}$ is distributed as uniformly as possible within $[\sigma_{\min}, \sigma_{\max}]$. To quantify this uniformity, we define the *spectrum entropy*

$$\mathcal{S} = -\int g(\sigma)\,\log g(\sigma)\,d\sigma,$$

where $g(\sigma)$ is the normalized spectral density. Maximizing $\mathcal{S}$ alone would distribute eigenvalues evenly, but functional sensitivity requires spectral weight to concentrate in the slow modes. Hence robustness and sensitivity impose opposing demands, which can be reconciled through the principle of maximum entropy with simultaneous constraints:

$$\hat{\mathcal{S}} = -\int g(\sigma)\,\log g(\sigma)\,d\sigma + \nu_0\Big(\int g(\sigma)\,d\sigma - 1\Big) + \nu_1\Big(\int \sigma g(\sigma)\,d\sigma - m\Big) + \mu\Big(\int \log\sigma\, g(\sigma)\,d\sigma\Big),$$

which is the standard form of constrained entropy maximization. Here, $\nu_0$ enforces normalization, $\nu_1$ fixes the mean eigenvalue, and $\mu$ incorporates the fluctuation entropy constraint. Taking the functional derivative with respect to $g(\sigma)$ yields

$$\frac{\delta\hat{\mathcal{S}}}{\delta g(\sigma)} = -\big(\ln g(\sigma) + 1\big) + \nu_0 + \nu_1\sigma + \mu\ln\sigma.$$

Setting this to zero gives

$$g^*(\sigma) = e^{-1+\nu_0+\nu_1\sigma}\,\sigma^{\mu},$$

a power-law form with an exponential cutoff. The exponent $\mu$ is defined at the single-protein level and reflects system size $N$ and structural features such as CO, so it does not map directly onto the global scaling exponent measured across ensembles. Nevertheless, the interpretation is consistent: $\mu$ characterizes how the stability–flexibility balance is resolved in an individual protein, while the global power-law scaling reflects the same critical mechanism across proteins.

This spectral perspective has a natural graph-theoretic interpretation. By the matrix–tree theorem, the fluctuation entropy $S$ is proportional to the logarithm of the number of spanning trees of the contact graph. For a fixed number of edges (i.e., fixed Laplacian trace), reducing the number of spanning trees increases $S$ and promotes power-law spectra. Evolutionarily, fewer spanning trees stabilize the hydrophobic core while enhancing collective flexibility, enabling proteins to tolerate mutations without losing functional responsiveness. Minimizing the number of spanning trees thus provides a structural mechanism by which robustness and sensitivity are jointly embedded into residue-contact topology.

### VI. TOPOLOGY–DYNAMICS RELATIONSHIPS BEYOND THE ENM APPROXIMATION

In this section, we show that the relationships between folding topology (CO), fluctuation entropy ($S$), and conformational flexibility extend beyond the harmonic ENM model, as supported by independent structural descriptors and experimental thermal stability data.

#### A. Structural Topology Measures: Modularity and Fractal Dimension

We examined two additional structural descriptors—network modularity $Q$ and fractal dimension $d_f$—to evaluate whether the topology–dynamics trends associated with CO and fluctuation entropy $S$ extend beyond the linear ENM approximation.

*a. Modularity $Q$.* Network modularity $Q$ measures the extent to which residues form densely connected substructures. It is defined as

$$Q = \frac{1}{2m}\sum_{i,j}\left[A_{ij} - \frac{k_i k_j}{2m}\right]\delta(c_i, c_j), \qquad \text{(S13)}$$

where $A_{ij}$ is the adjacency matrix, $k_i = \sum_j A_{ij}$ is the degree of node $i$, $m = \frac{1}{2}\sum_{i,j} A_{ij}$ is the total number of edges, and $c_i$ is the module assignment of residue $i$. We computed $Q$ on residue contact networks constructed with a cutoff of $r_c = 8$ Å and determined community structure using the Louvain method.

*b. Fractal dimension $d_f$.* To quantify packing density, we estimated the fractal dimension $d_f$ using a scaling relation of the form $n(r) \sim r^{d_f}$, where $n(r)$ is the number of residues within distance $r$ of a reference residue. For each protein, $d_f$ was computed across multiple reference points and averaged.

*c. Results.* As shown in Fig. S13, $Q$ correlates negatively with CO and positively with $S$ for proteins of similar length. Conversely, $d_f$ correlates positively with CO and negatively with $S$. Thus, proteins with more sequence-local contacts tend to form more modular and less densely packed structures, whereas proteins with more long-range contacts exhibit denser packing and reduced flexibility.

These consistent relationships show that the topology–dynamics trend captured by CO and $S$ does not arise from the linear harmonic assumptions of the ENM. Both modularity $Q$ and fractal dimension $d_f$ are intrinsically nonlinear geometric descriptors: $Q$ characterizes the combinatorial organization of residue communities, while $d_f$ quantifies multiscale packing density via power-law scaling. Their agreement with CO and $S$ indicates that the observed trend reflects underlying geometric constraints of folded polymers, specifically how contacts are distributed across length scales, rather than any particular linear mode approximation. Physically, the CO–$S$ relationship therefore encodes global topological con-

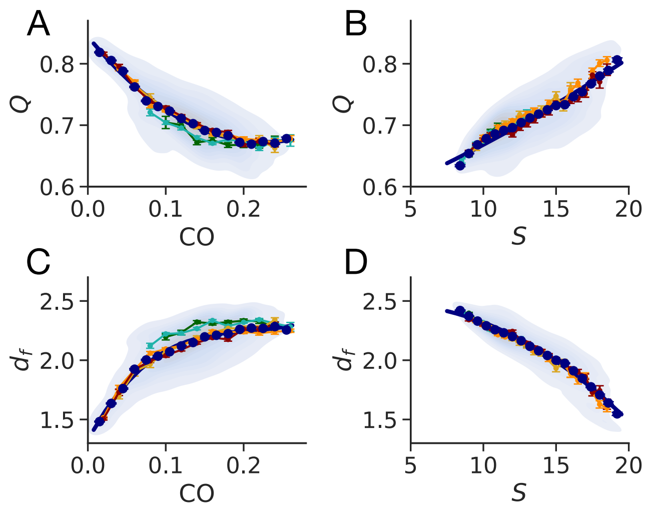


FIG. S13. Relationships between network modularity ($Q$), fractal dimension ($d_f$), contact order (CO), and fluctuation entropy ($S$). (A) $Q$ vs. CO, (B) $Q$ vs. $S$, (C) $d_f$ vs. CO, and (D) $d_f$ vs. $S$. Shaded regions show data density; lines indicate average trends. Color scheme matches Main Text Fig. 2.

straints that limit how a chain can pack and fluctuate, independent of the modeling framework.

#### B. Consistency with Experimental Melting Temperatures

To evaluate whether the CO–$S$ relationship reflects intrinsic thermodynamic stability, we compared CO and $S$ with melting temperatures ($T_m$) from the Meltome Atlas [21]. For each protein, we assigned a single representative $T_m$ by taking the median across all reported measurements. We mapped Meltome gene identifiers to UniProt accessions, retrieved the corresponding AF2 structures, and retained only one-to-one mappings. CO and $S$ were computed as described in the main text. We focused on a single species because thermostability is shaped mainly by environmental pressures and evolutionary selection rather than by organismal complexity; cross-species comparisons would therefore add unnecessary heterogeneity. The Meltome Atlas provides the densest measurements for human proteins, so we restricted the analysis to the human proteome.

In addition to the chain-length group analyzed in the main text ($225 \leq N < 275$), we examined proteins with $300 \leq N < 400$ and $400 \leq N < 500$ (Fig. S14A–D). Across all size ranges, CO increases with $T_m$ and $S$ decreases with $T_m$, with median trends closely matching full-data fits, indicating robustness to sampling variation.

Together, the correspondence between CO, $S$, and experimentally measured $T_m$ shows that the CO–$S$ trend reflects intrinsic biophysical constraints on protein stability. CO and $S$ thus provide experimentally grounded, physically interpretable descriptors linking topology, native-state flexibility, and thermo-stability.

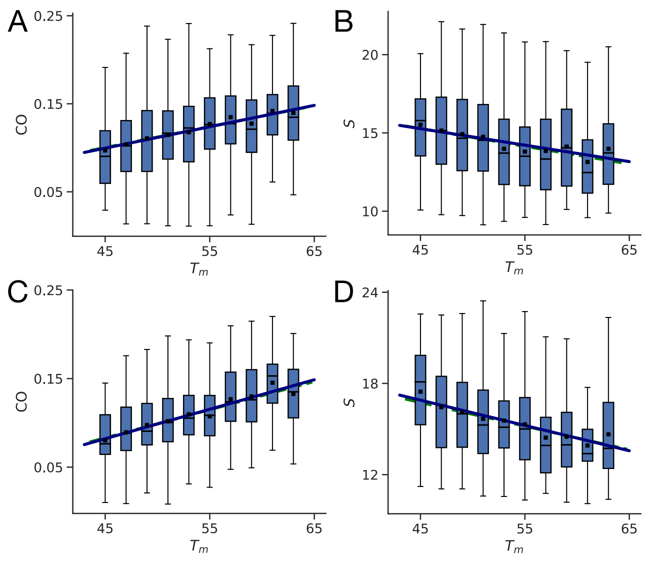


FIG. S14. Thermal stability validation of the CO–$S$ relation for human proteins of different chain lengths. For proteins with $300 \leq N < 400$ (A,B) and $400 \leq N < 500$ (C,D), CO increases with melting temperature $T_m$ (A,C), and the corresponding $S$ decreases with $T_m$ (B,D). In all the panels, fits to the full data (dashed lines) and to the binned means (dots with solid-line fits) nearly overlap, indicating consistent trends.

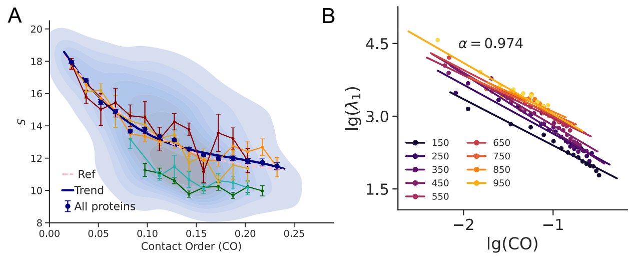


FIG. S15. Robustness to sequence redundancy. Using a 50%-identity redundancy-reduced dataset (sequences clustered at 50% identity using MMseqs2), we repeated the CO–$S$ analysis for proteins of similar chain length ($N \approx 250$). (A) The CO–$S$ relationship agrees closely with the full dataset (pink line; Fig. 2A). (B) The scaling analysis yields fitted exponents consistent with those obtained from the complete proteome.

### VII. THE ROBUSTNESS OF SCALING ANALYSIS AND RENORMALIZATION

#### A. Robustness to sequence redundancy

To test whether the CO–$S$ relationship is affected by sequence redundancy, we constructed a redundancy-reduced dataset by clustering proteins with more than

50% sequence identity using MMseqs2 (minimum identity 50%, at least 80% alignment coverage) and retaining one representative per cluster; an additional analysis at 30% identity gave identical results. As shown in Fig. S15A, the CO–$S$ relationship closely matches that of the full dataset (pink line in Fig. 2A), with slightly larger fluctuations due to reduced sample size, while the corresponding scaling analysis yields nearly identical exponents (Fig. S15B). Thus, redundancy filtering removes close homologs without altering the CO–$S$ trend or fitted scaling parameters, confirming that the observed relationships are not artifacts of sequence redundancy.

#### B. Data binning and fitting robustness

In this subsection, we systematically varied the binning parameters and compared the resulting scaling fits. We first examined the scaling exponent $\alpha$ (main text Fig. 3A) under three controlled perturbations to the binning scheme (Fig. S16): (A) an *N-shift test*, where the bin width in $N$ was fixed but the starting point of $N$ intervals was varied; (B) an *N-resolution test*, where bin widths in $N$ were changed while keeping the starting point fixed; and (C) a *CO-resolution test*, where the number of CO bins was varied while keeping $N$ binning fixed. In all cases, the fitted exponents and overall scaling trends remained stable, indicating robustness to bin alignment, $N$ resolution, and CO resolution.

We also tested the robustness of the scaling exponent $\zeta$ (main text Fig. 3D) across different binning strategies using the same set of proteins (Fig. S17). Variations included: (A–C) different bin widths for $N$ (80, 150, and 200; main text uses 100); (D–F) different starting points for $N$ bins while keeping the bin width at 100 (e.g., starting at 25 corresponds to $125 \leq N < 225$, starting at 50 to $250 \leq N < 350$, etc.). These tests yielded highly consistent $\zeta$ estimates, with a mean of 2.352 and a standard deviation of 0.045. The binning-robustness tests demonstrate that, within reasonable parameter ranges, our conclusions are stable with respect to binning choices.

#### C. Secondary-structure composition

In this subsection, we assess whether the scaling relations identified in the main text remain valid across proteins with different secondary-structure compositions, used here as a practical proxy for detecting potential bias arising from the overrepresentation of certain structural classes. Because complete fold annotations are not available for all proteins in the dataset, and since folds are defined at the domain rather than the whole-protein level, we instead use secondary-structure content as a uniformly applicable descriptor at the whole-protein scale. Using DSSP [22], we calculated the $\alpha$-helix fraction $f_\alpha$ and $\beta$-sheet fraction $f_\beta$ for each protein, with the turn/coil fraction given by $f_{\rm tc} = 1 - f_\alpha - f_\beta$. Proteins

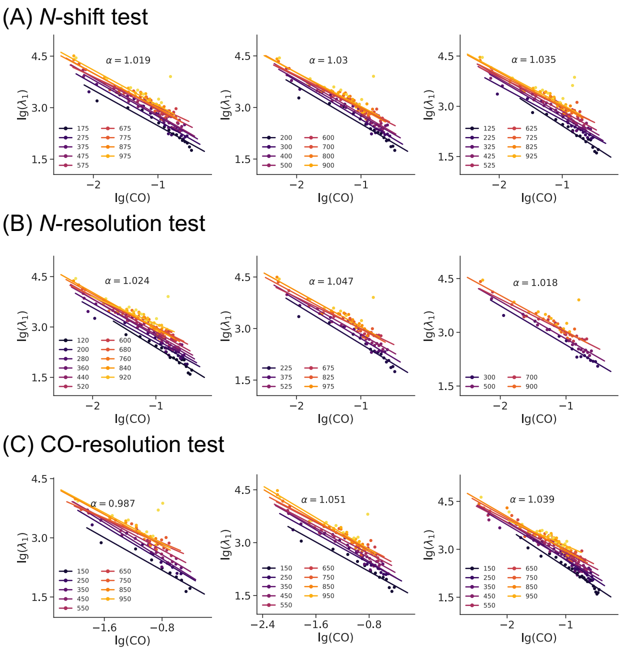


FIG. S16. Robustness of scaling results to binning setups. (A) *N-shift test*: same bin width in $N$ but varying the starting point of $N$ bin intervals, CO bin resolution fixed. (B) *N-resolution test*: different bin widths in $N$ with the same starting point, CO bin resolution fixed. (C) *CO-resolution test*: same $N$ binning as in the main text but varying the number of CO bins. All tests confirm that the fitted exponents are insensitive to bin alignment $N$ and CO resolution.

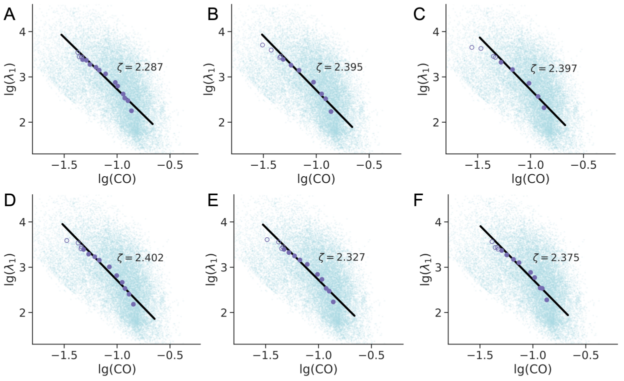


FIG. S17. Binning-robustness test for $\zeta$. Fitting of $\zeta$ using the same group of proteins under different binning setups. (A–C) Different bin widths for $N$: (A) 80, (B) 150, and (C) 200 (main text uses 100). (D–F) Different bin starting points with bin width 100: (D) start at 25, (E) start at 50, and (F) start at 75. For example, starting at 25 corresponds to $125 \leq N < 225$, starting at 50 to $250 \leq N < 350$, etc.

were then classified into four broad composition groups based on $(f_\alpha, f_\beta)$ thresholds (Fig. S18A):

- *Helix-rich*: $f_\alpha + f_\beta \geq 0.5$ and $f_\alpha - f_\beta > 0.2$
- *Sheet-rich*: $f_\alpha + f_\beta \geq 0.5$ and $f_\beta - f_\alpha > 0.2$

- *Helix/sheet-balanced*: $f_\alpha + f_\beta \geq 0.5$ and $|f_\alpha - f_\beta| \leq 0.2$
- *Turn/coil-rich*: $f_\alpha + f_\beta < 0.5$

While this approach is not equivalent to a fold-specific analysis, it provides a tractable way to examine whether the observed scaling behavior is preserved across distinct broad-scale structural architectures. These groups are shown in Fig. S18A, with their CO and $S$ distributions in Fig. S18B–C. Composition effects are evident: turn/coil–rich proteins tend to have lower CO and higher $S$, while helix-rich proteins have slightly lower CO and higher $S$ than sheet-rich or balanced proteins.

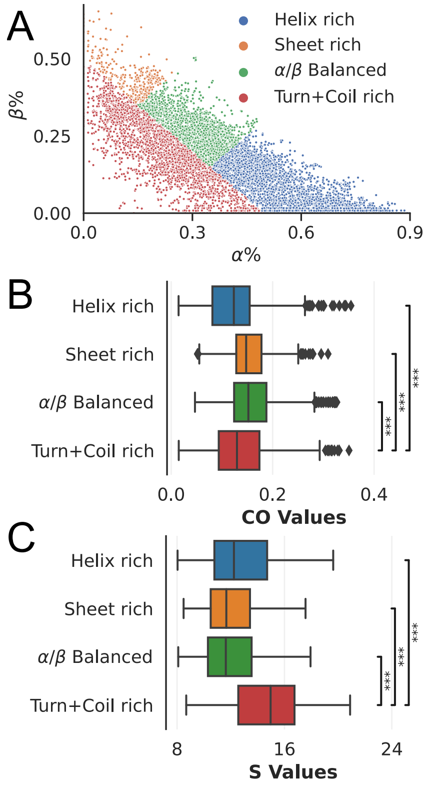


FIG. S18. Structural composition effects. (A) Classification of proteins into four secondary-structure composition groups in the $(f_\alpha, f_\beta)$ plane: helix-rich (red), sheet-rich (blue), helix/sheet-balanced (green), and turn/coil-rich (orange). (B) CO and (C) $S$ distributions by groups.

Despite these composition differences, all four groups preserve the same power-law scaling form, although the fitted exponents $(\alpha, \beta, \gamma, \eta)$ differ across groups. For illustration, we focus on the largest group—helix-rich proteins—which provides the best statistics. The fits (Fig. S19) yield $\alpha = 1.067$, $\beta = 1.631$, $\gamma = 0.99$, and $\eta = 0.62$, satisfying $\alpha\eta \approx \beta - \gamma$ $(0.66 \approx 0.64)$. This indicates that while structural composition bias can shift absolute exponent values, the internal consistency of the scaling laws is maintained.

Secondary-structure composition analysis confirms that the observed scaling relations are preserved across broad architectural categories. While composition can influence absolute exponent values, the scaling form and internal consistency relations remain valid, indicating that our findings are not driven by overrepresentation of a particular structural class.

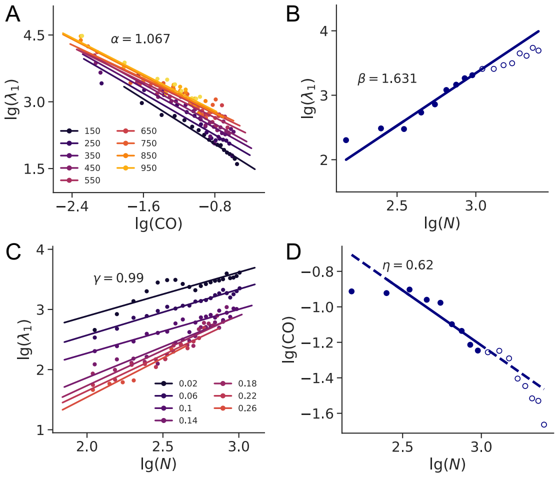


FIG. S19. Scaling fits for helix-rich proteins: (A) $\alpha$, (B) $\beta$, (C) $\gamma$, (D) $\eta$. Aside from selecting helix-rich proteins, fitting procedures are identical to those in Fig. 3 of the main text.

### D. Robustness of Scaling Across Organisms and Mode Numbers

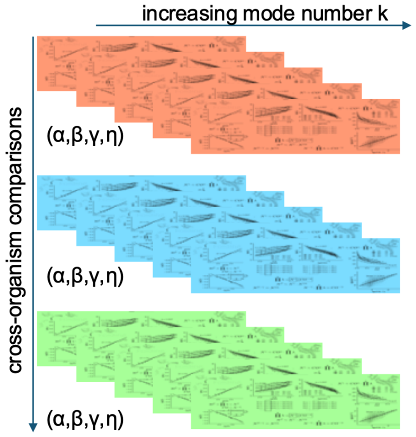


FIG. S20. Schematic illustrating validation of the scaling relation across organisms (vertical axis) and number of slow modes considered (horizontal axis).

To assess the robustness of the proposed scaling law, we repeated our analysis across individual model organisms and across varying numbers of slow modes. Specifically, to test organismal dependence, we applied the scaling analysis to proteins from representative species such as *E. coli*, yeast, and human (vertical axis in Fig. S20).

In each case, we consistently observed a negative correlation between CO and the leading eigenvalue $\lambda_1$, the power-law scaling relation $\lambda_1 \sim \mathrm{CO}^{-\alpha}$, and the validity of the scaling relation $\alpha\eta = \beta - \gamma$.

We further examined the impact of the number of slow modes by replacing $\lambda_1$ with the cumulative product $\prod_{t=1}^{k} \lambda_t$ for $k = 1, 2, \ldots, 10$ (horizontal axis in Fig. S20). For each $k$, we recalculated the scaling exponents $\alpha^{(k)}, \beta^{(k)}, \gamma^{(k)}$ using the same linear regression procedure on log-log transformed data as in the main text. The resulting values satisfy the relation $\eta = (\beta^{(k)} - \gamma^{(k)})/\alpha^{(k)}$, with $\eta$ remaining stable across different $k$. Since $\log \prod_{t=1}^{k} \lambda_t = \sum_{t=1}^{k} \log \lambda_t$, this corresponds directly to the truncated fluctuation entropy $S_k$.

The fitted exponents $\alpha^{(k)}, \beta^{(k)}, \gamma^{(k)}$ for representative values $k = 2, 4, 6, 8, 10$ are shown in Fig. S21, using the same protein dataset as in Main Text Fig. 3. Each row corresponds to a different $k$, and each column to a different exponent. As $S_k$ increases with $k$, the exponent values shift accordingly, yet the scaling relation involving $\eta$ consistently holds. For example, for $k = 2$, we obtain $\alpha = 1.959$, $\beta = 3.073$, and $\gamma = 1.698$, yielding $\eta \approx 0.654$, which closely matches the directly fitted value $\eta = 0.681$ (Main Text Fig. 3E, inset). Similar consistency is observed when the analysis is repeated for proteins from individual organisms: although exponent values may vary across species and $k$, the underlying scaling relation remains robust (Main Text Fig. 3F).

#### E. Renormalization Analysis

To test whether the observed scaling reflects physical constraints rather than biological trends, we performed a data-driven renormalization analysis. Specifically, we applied structural coarse-graining to assess the persistence of the CO–$S$ relationship across resolutions.

The first strategy mimics real-space renormalization. We selected 500 proteins from the dataset and systematically grouped sequence-adjacent residues into coarse-grained units—e.g., 2-to-1, 3-to-1, up to 10-to-1—and constructed a new contact map at each resolution. Two coarse-grained "residues" were considered in contact if at least one pair of their constituent residues was in contact at the finer scale. At the base level, contacts were defined as in the Main Text: Residue pairs with $|i - j| \geq 4$ and spatial distance less than $r_{\mathrm{CO}}$. For each coarse-grained structure, we computed the corresponding CO and fluctuation entropy $S$, with $S$ approximated by the logarithm of the leading Laplacian eigenvalue, $\lambda_1$. This process yielded a sequence of $(\mathrm{CO}, \lambda_1)$ pairs across renormalization levels (Fig. S22A).

The second strategy samples structural segments from longer proteins, treating each fragment as an independent "subprotein." Using the same set of 500 proteins, we randomly selected segments of length 1/2, 1/3, ..., 1/10 of the full chain, sampling each size 20 times by choosing random starting positions along the sequence. For each

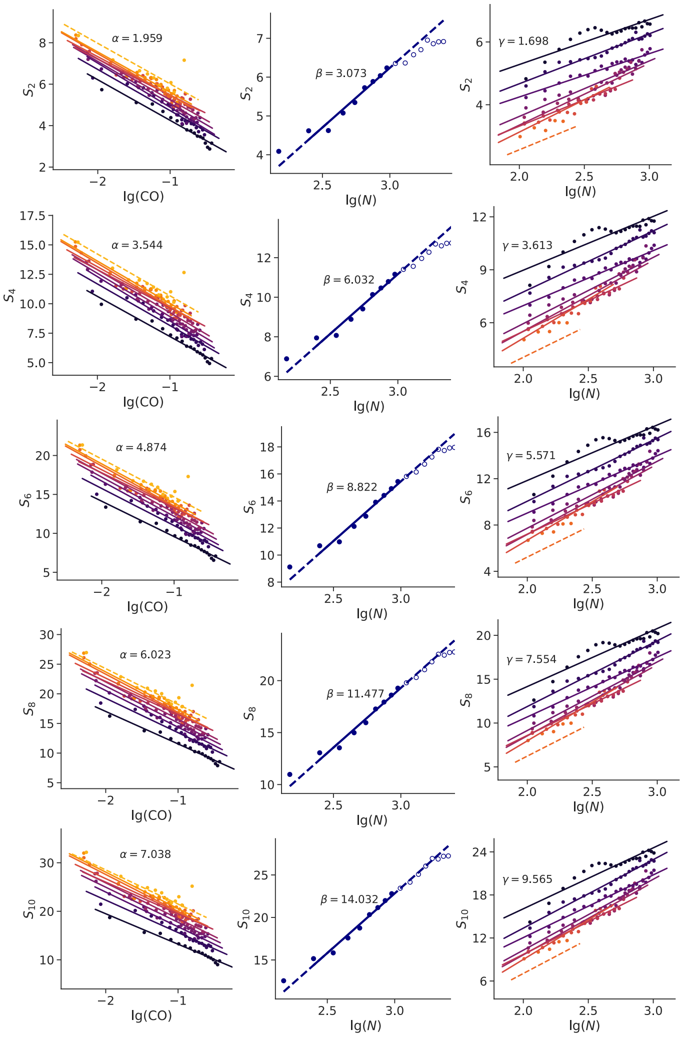


FIG. S21. Fitted scaling exponents $\alpha^{(k)}$, $\beta^{(k)}$, and $\gamma^{(k)}$ for different numbers of slow modes $k = 2, 4, 6, 8, 10$, using the same dataset as Main Text Fig. 3. Each row corresponds to a different value of $k$, and each column shows the respective exponent. Despite variations in the fitted values, the relation $\eta = (\beta^{(k)} - \gamma^{(k)})/\alpha^{(k)}$ holds across all cases.

subprotein, we computed CO and $\lambda_1$ using the same definitions and aggregated the results across all samples.

In both strategies, the log-log plots of $\lambda_1$ versus CO exhibit consistent power-law behavior, closely aligning with the global scaling relation $\lambda_1 \sim \mathrm{CO}^{-\zeta}$ observed in the Main Text (Fig. 3D). As shown in Fig. S22B, most renormalized or sampled structures fall along a reference line with slope $-\zeta$, where $\zeta = 2.293$ was obtained from proteome-wide analysis (Fig. 3D). This result demonstrates that the scaling behavior is not exclusive to naturally evolved proteins, but also holds across systematically coarse-grained and fragmented structures. Such invariance under resolution and size reduction strongly supports the physical origin of the observed scaling, analogous to renormalization flow in statistical physics.

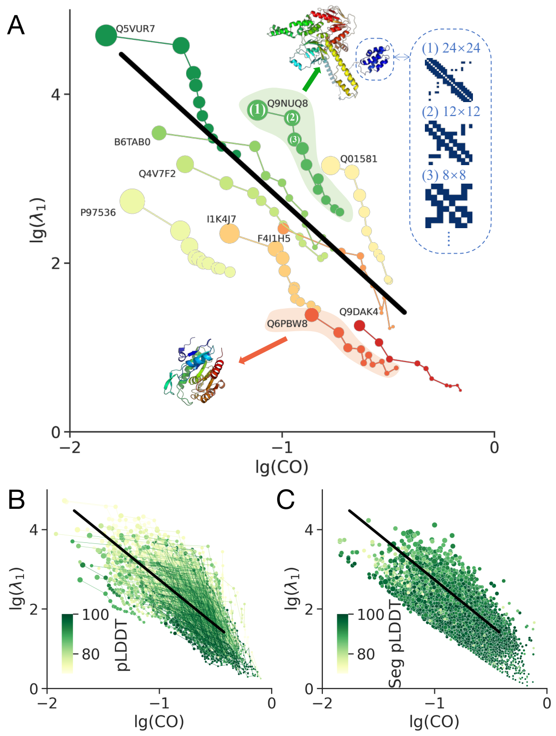


FIG. S22. Renormalization analysis reveals the physical invariance of the scaling. (A) Schematic of hierarchical coarse-graining applied to protein contact maps. Each level corresponds to grouping residues into larger coarse-grained units, and computing contact order (CO) and fluctuation entropy (approximated by $\log \lambda_1$) for each resolution. (B-C) Log-log plots of $\lambda_1$ versus CO for (B) renormalized structures and (C) sampled segments. The black lines show the reference slope $-\zeta = -2.293$ from proteome-wide fits (Fig. 3D). Both coarse-grained proteins and structural fragments align with the expected scaling, highlighting the robustness and physical origin of the observed relation.

### VIII. POLYMER-PHYSICS ORIGIN OF SCALING EXPONENTS

In Sec. V, we showed that fluctuation entropy $S$ is directly linked to the number of spanning trees in the residue contact network, with maximizing $S$ (equivalently, minimizing the number of spanning trees) shaping the vibrational spectrum. While this identifies a general optimization principle, it does not specify the physical interactions that realize these constraints. Polymer theory provides this physical baseline: in compact globular proteins, contact formation follows loop-closure statistics dictated by chain connectivity and excluded-volume effects, establishing the structural scaffold on which evolutionary and functional pressures act.

Let $P(s)$ denote the probability that two residues separated by a sequence distance $s = |i - j|$ form a contact. In compact chains, loop-closure statistics and steric constraints yield [23–26]

$$P(s) \propto s^{-c}, \quad s_0 \le s \le s_*(N), \quad c > 1.$$

For an ideal Gaussian chain in three dimensions, random-walk statistics give the mean spatial separation

$$R(s) \equiv \langle |\mathbf{r}_i - \mathbf{r}_j| \rangle \sim s^{1/2},$$

and the contact probability scales as the inverse accessible volume,

$$P(s) \propto [R(s)]^{-3} \propto s^{-3/2},$$

giving the classic Gaussian-chain result $c = 3/2$. Here, $s_0 = O(1)$ is the minimal loop length set by local backbone geometry and excluded volume—below this scale, residues cannot fluctuate independently. The upper cutoff $s_*(N)$ is the largest effective loop length, limited by finite chain size or modular/domain boundaries.

Empirically, the mean per-residue contact degree is nearly size-independent, so contact-number effects cancel in the mean separation:

$$\langle s \rangle = \frac{\int_{s_0}^{s_*} s P(s)\, ds}{\int_{s_0}^{s_*} P(s)\, ds} = \frac{\int_{s_0}^{s_*} s^{1-c}\, ds}{\int_{s_0}^{s_*} s^{-c}\, ds}.$$

The asymptotic behavior of this ratio depends on the loop-closure exponent $c$:

*(i) $c > 2$ (narrow loop-length spectrum):*

$$\langle s \rangle \to \frac{c-1}{c-2} s_0 = O(1), \quad \mathrm{CO} \sim N^{-1} \quad \Rightarrow \quad \eta = 1.$$

*(ii) $c = 2$ (marginal spectrum):*

$$\langle s \rangle \sim s_0 \log \frac{s_*}{s_0}, \quad \mathrm{CO} \sim \frac{\log N}{N},$$

giving $\eta \approx 1$ (with logarithmic corrections).

*(iii) $1 < c < 2$ (broad loop-length spectrum):* Here, the largest accessible loops dominate the numerator of the contact-order integral, while the shortest loops control the denominator:

$$\langle s \rangle \sim \frac{c-1}{2-c} s_0^{c-1} s_*^{2-c}.$$

If $s_*(N) \sim N^{\phi}$ with $0 < \phi < 1$, then

$$\mathrm{CO} = \frac{\langle s \rangle}{N} \sim N^{(2-c)\phi - 1} \quad \Rightarrow \quad \eta = 1 - (2-c)\phi.$$

For typical dataset parameters ($N \approx 250$, $s_0 \sim 1$–$3$), we have $s_*/s_0 \gtrsim 50$, ensuring a broad enough scaling window for these asymptotics to apply. The condition $(s_*/s_0)^{2-c} \gg 1$ is also well satisfied.

Polymer statistics provide a general baseline:

$$\eta = 1 - (2-c)\phi, \quad 0 < \phi < 1,$$

where $c$ reflects loop-closure geometry and $\phi$ the scaling of the largest effective loop with chain size. From a criticality perspective, these parameters set the balance between local stability from short loops and global flexibility from large loops, thereby shaping the spectrum of slow collective modes.

This polymer-scaling framework shall be viewed as a heuristic ansatz: the $c = 3/2$ exponent holds exactly only for ideal Gaussian chains, whereas real proteins are subject to additional geometric, packing, and evolutionary constraints. In practice, physical couplings, mutational constraints, coevolutionary signals, and functional demands can shift $(c, \phi)$ toward a balance characteristic of near-critical systems—enhancing long-range correlations, extending residue–residue coupling ranges, and modifying $P(s)$ and $s_*(N)$ beyond the polymer limit while preserving the overall scaling form.

The empirical relation $\alpha\,\eta = \beta - \gamma$ can thus be viewed as a *consistency condition* arising from separability and the observed CO $\sim N^{-\eta}$. Loop-closure geometry and chain topology set the physical baseline; evolutionary and functional constraints refine it. Identifying statistical signatures of these refinements in modern AI protein models, and linking them to their physical–evolutionary origins, remains an important open question.

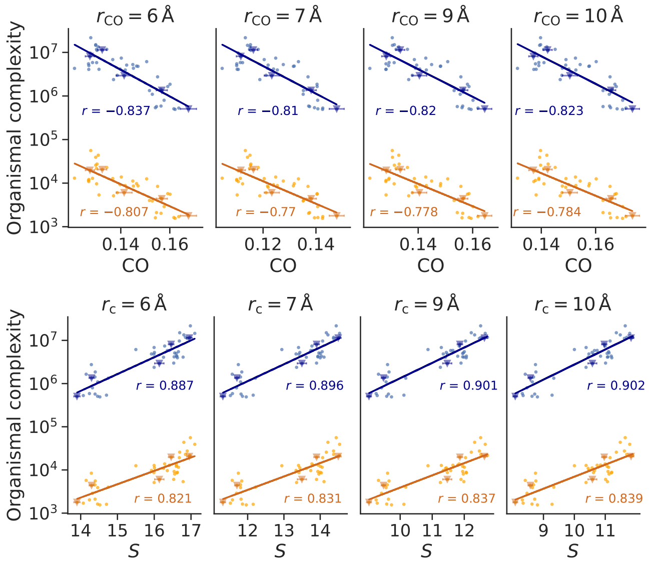


FIG. S23. Validation of Main Text Figs. 4A&B with residue-contact cutoffs ($r_{CO} = 6, 7, 9,$ and 10 Å). The negative correlation between organismal complexity and mean CO (top) and the positive correlation with $S$ (bottom) remain robust. Blue points and fits use complexity measured by total proteome chain length; orange uses the number of distinct proteins. Pearson $r$-values are reported for each fit. Five organisms (*M. jannaschii*, *E. coli*, *S. cerevisiae*, *C. elegans*, and *H. sapiens*) are marked with inverted triangles. Proteins with $225 \le N < 275$ and mean pLDDT $\ge 70$ are used.

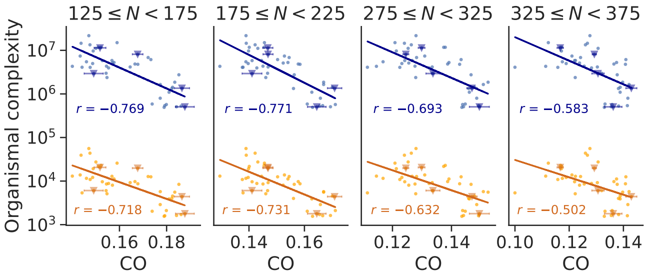


FIG. S24. Validation of Main Text Fig. 4A using different chain-length groups ($N_g = 150, 200, 300,$ and 350), where each group includes proteins with $N_g - 25 \le N < N_g + 25$ and mean pLDDT $\ge 70$. Results confirm the robustness of the negative correlation between organismal complexity and mean CO. The residue contact cutoff is $r_{CO} = 8$ Å; all other settings match Fig. S23.

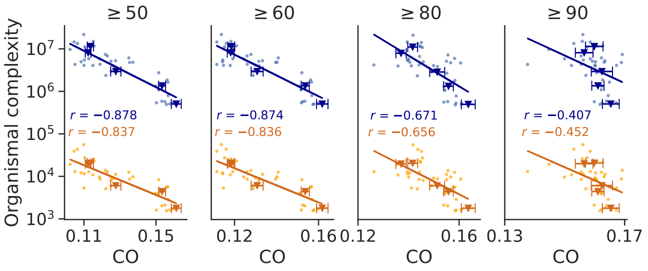


FIG. S25. Validation of Main Text Fig. 4A using different pLDDT thresholds ($\ge$ 50, 60, 80, and 90) to test robustness of the chosen AF2 confidence cutoff. As in the Main Text, proteins of similar length ($225 \le N < 275$) and a contact cutoff of $r_{CO} = 8$ Å are used. All other settings follow Fig. S23.

### IX. ROBUSTNESS OF CROSS-ORGANISM ANALYSIS

To confirm the robustness of the organismal complexity trends reported in Fig. 4A and 4B, we analyzed proteins from 45 organisms under varying residue-contact definitions (distance cutoffs $r_{CO}$ from 6 to 10 Å), different chain-length windows, and alternative protein-filtering thresholds. As shown in Fig. S23, the negative correlation between organismal complexity and mean CO, and the positive correlation with fluctuation entropy $S$, remain consistent across all conditions.

As shown in Fig. S24, the negative correlation between mean CO and organismal complexity remains robust across different protein chain length groups ($N_g$ = 150, 200, 300, and 350). Across all groups, the negative correlation persists, demonstrating that the observed relationship is not sensitive to the specific choice of chain length window. These results confirm that the correlation between organismal complexity and CO is a robust statistical feature across proteins of varying lengths.

As shown in Fig. S25, varying the pLDDT threshold ($\ge$ 50, 60, 80, and 90) does not substantially affect the observed negative correlation between mean CO and organismal complexity. However, applying very stringent pLDDT cutoffs (e.g., $> 90$) may introduce bias by preferentially selecting highly ordered proteins that lack flexible linkers or disordered regions, narrowing the structural diversity toward more stable conformations. Despite this potential bias, the negative correlation remains statistically significant even under the highest threshold.

[15] G. Kirchhoff, Ueber die auflösung der gleichungen, auf welche man bei der untersuchung der linearen vertheilung galvanischer ströme geführt wird, Ann. Phys. **148**, 497 (1847).

[16] S. Chaiken and D. J. Kleitman, Matrix tree theorems, J. Combin. Theory A **24**, 377 (1978).

[17] R. B. Bapat, *Graphs and Matrices* (Springer, London, 2010).

[23] P.-G. de Gennes, *Scaling Concepts in Polymer Physics* (Cornell University Press, Ithaca, NY, 1979).

[24] M. Doi and S. F. Edwards, *The Theory of Polymer Dynamics* (Oxford University Press, Oxford, 1986).

[25] J. des Cloizeaux, Short range correlation between elements of a long polymer in a good solvent, Journal de Physique **41**, 223 (1980).

[26] J. Shimada and H. Yamakawa, Ring-closure probabilities for twisted wormlike chains: Application to DNA, Macromolecules **17**, 689 (1984).
